# Active zone maturation state controls synaptic output and release mode and is differentially regulated by neuronal activity

**DOI:** 10.1101/2025.02.03.636302

**Authors:** Yulia Akbergenova, Jessica Matthias, J. Troy Littleton

**Affiliations:** The Picower Institute for Learning and Memory, Department of Brain and Cognitive Sciences, Department of Biology, Massachusetts Institute of Technology, Cambridge, MA 02139; Abberior Instruments America, Bethesda, MD

## Abstract

Synapse formation requires the gradual accumulation of cytomatrix proteins and voltage-gated Ca^2+^ channels (VDCCs) at presynaptic active zones (AZs) to support neurotransmitter release. To correlate AZ maturation with synaptic output, quantal imaging was performed at serially imaged time-stamped *Drosophila* synapses. Evoked release strength correlated strongly with AZ age and accumulation of late AZ scaffolds, while immature sites lacking VDCC accumulation supported spontaneous release. To examine how neuronal activity regulates AZ maturation and protein accumulation, the effects of disruptions to SV fusion or action potential generation were analyzed. Decreasing neuronal activity reduced AZ seeding and caused hyperaccumulation of presynaptic material at existing AZs. Although enlarged AZs are also observed in *rab3* mutants, activity reduction acted through an independent mechanism that required postsynaptic glutamate receptor-dependent signaling. Together, these data indicate AZ maturation state sets distinct presynaptic release modes and output strength, with neuronal activity shaping both AZ number and size across development.

## Introduction

Synaptogenesis requires the formation of highly specialized points of contact between pre and postsynaptic cells. Active zones (AZ) specialized for neurotransmitter release assemble at sites along the presynaptic axon opposed to neurotransmitter receptor clusters from postsynaptic partners^1^. AZs form an electron-dense cytomatrix containing multiple evolutionary conserved proteins, including cell adhesion molecules (Neurexins, Teneurins), scaffolding proteins (Liprin-α, Syd-1, RIM, RIM-Binding Protein (RBP), Unc13, BRP/ELKS/CAST) and Ca_v_2 voltage-gated Ca^2+^ channels (VDCCs)^2,3^. A set of early AZ scaffolding proteins initially assemble at release sites before the arrival of VDCCs and late scaffolds that cluster synaptic vesicles (SVs)^3^. Although most AZs contain a similar assortment of proteins, the distribution of these components is not uniform and can be rapidly modified^4–9^. How the differential distribution of specific AZ components affects synaptic output is not clear. Similarly, mechanisms that shape presynaptic AZ assembly and remodeling in response to changes in neuronal activity remain elusive.

*Drosophila* larval glutamatergic neuromuscular junctions (NMJs) have emerged as a useful model connection to characterize AZ maturation and synaptic output. NMJs form during late embryogenesis and expand throughout the subsequent 6 days of larval development^2^. Motoneurons (MNs) continue to add new synaptic boutons that each contain many AZs, with Bone Morphogenetic Protein (BMP) and Wingless signaling among others, coordinating NMJ growth^10^. AZ maturation proceeds through the accumulation of molecular components that can be resolved with live imaging through the cuticle using fluorescently-tagged proteins of interest^7,11^. With the advent of quantal imaging that allows SV release to be measured at individual release sites^12,13^, one can determine how the specific molecular composition of AZs correlates with whether they release only spontaneous SVs, have low evoked release probability (*P_r_*) or instead function as high *P_r_* sites. In the current study, we examine how AZ material accumulation regulates synaptic output and how this process is modulated by alterations in MN activity.

By timestamping synapses with a postsynaptic glutamate receptor (GluR) subunit linked to the photoconvertible fluorescent marker mMaple, we observe that presynaptic output dramatically increases as AZs mature over the course of several days. Although early scaffolding components such as Liprin-α, Syd1 and Unc13B are the first to arrive at developing AZs, their overall abundance correlates poorly with VDCC accumulation and the strength of evoked output at single release sites. In contrast, the abundance of late arriving scaffolding components like RBP, BRP and Unc13A, along with Cacophony (Cac, the *Drosophila* Ca_v_2 homolog), have a stronger contribution to synaptic efficacy. Maturing AZs that have not yet accumulated Cac channels appear as spontaneous-only sites that lack evoked release. The normal process of AZ formation and maturation is altered when action potential generation or SV release is disrupted in individual MNs. Instead of seeding new AZs throughout development, silenced neurons form fewer and larger release sites. AZ enlargement following disrupted presynaptic output requires postsynaptic GluRIIA function and acts through an independent mechanism from *rab3* mutants that also contain larger AZs. In contrast, enhanced activity leads to faster material accumulation but also sets an upper limit on synapse size. These data indicate neuronal activity regulates AZ seeding and material accumulation, with increases in functional output over time. Inactivity reduces AZ formation and increases AZ size, with AZ maturation occurring over a longer timescale and culminating in excess material at fewer release sites.

## Results

### A genetic toolkit for timestamping synapse age

Prior work using serial imaging of NMJ growth in briefly anesthetized *Drosophila* larvae suggested newly formed AZs are weaker than their more mature counterparts^7^. However, the rapid rate of AZ addition during larval growth makes it difficult to precisely follow defined AZs across multiple imaging sessions. To unambiguously determine how synapse age regulates presynaptic release mode and synaptic strength, a new toolkit for timestamping individual release sites was developed. mMaple is a genetically encoded fluorophore that undergoes complete green-to-red photoconversion (PC) upon illumination with 405 nm light^14^. By attaching mMaple to the core postsynaptic glutamate receptor subunit GluRIIE in transgenic animals, all postsynaptic densities (PSDs) opposed to presynaptic AZs at the NMJ can be reliably labeled (Fig. 1A). Brief 15 second illumination at 405 nm through the cuticle of intact 1^st^ instar larvae was sufficient to generate complete transition of the entire pool of GluRIIE^Maple^ from green to red at muscle 4 (M4) NMJs (Fig. 1B). Serial imaging of NMJs identified a small pool of non-synaptic red^+^ GluRIIE^Maple^ that continued to incorporate into PSDs during the 1^st^ day after PC, resulting in a mild increase in the number of red^+^ GluRIIE positive PSDs (Fig. 1C). After 24 hours, all newly formed PSDs contained only non-PC green^+^ GluRIIE^Maple^ (hereafter referred to as GluR^New^, Fig. 1C, D). Quantification of the number of red^+^ GluRIIE^MaplePHC^ positive PSDs (hereafter referred to as GluR^Old^) revealed no change between 24 and 48 hours (Fig. 1D), indicating the entire pool of previously PC GluRIIE^Maple^ had inserted into PSDs within a day and did not undergo lateral movement into newly formed synapses.

**Fig. 1.**
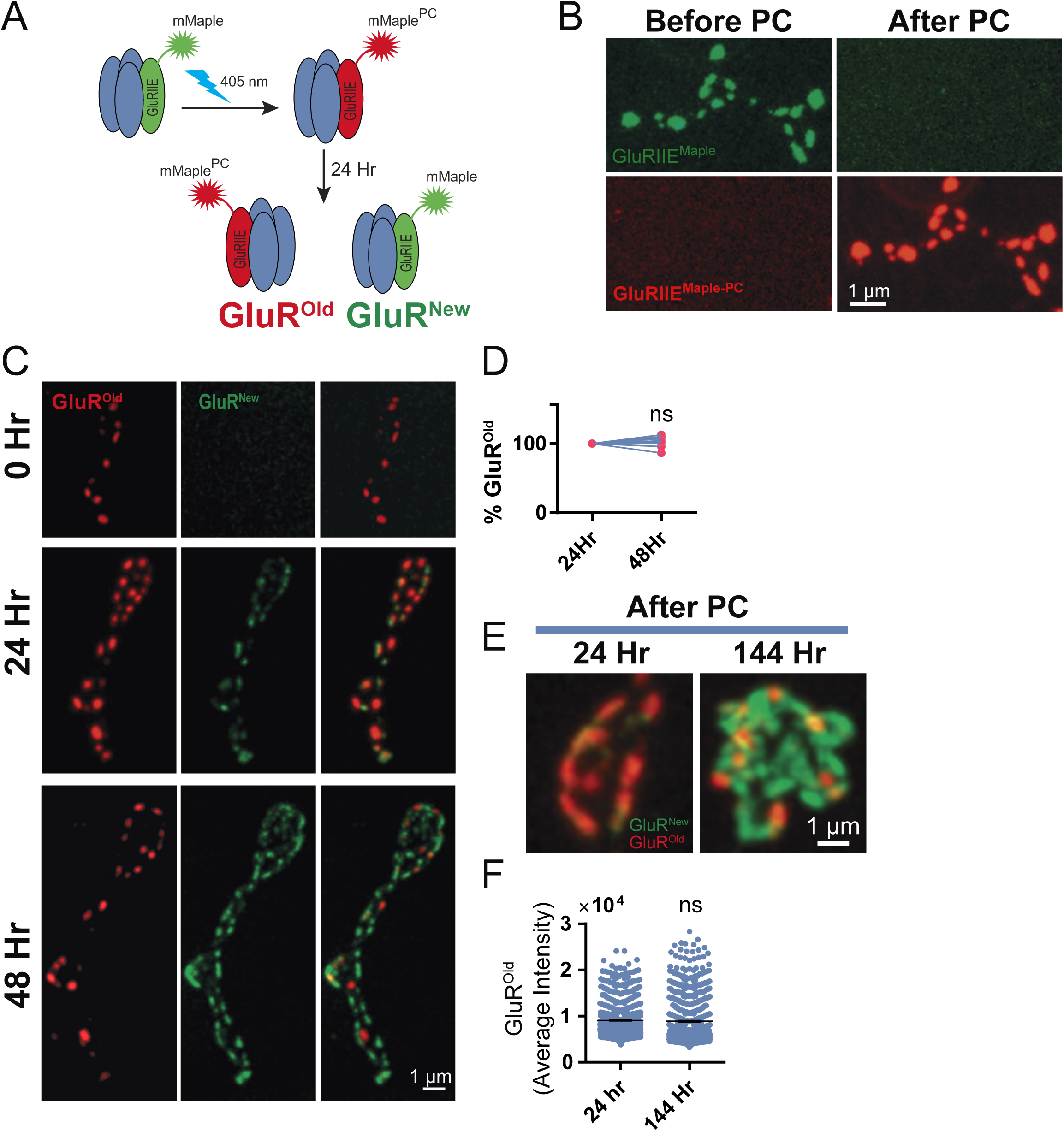
GluRIIE-mMaple photoconversion to timestamp synapse age. **A.** Schematic showing photoconversion (PC) of GluRIIE^Maple^ from green to red. All existing PSDs at the time of PC contain red^+^ GluRIIE (PSD^Old^), while newer PSDs formed after PC contain green^+^/red^-^ GluRIIE (PSD^New^). **B.** Complete PC of larval NMJ PSDs containing GluRIIE^Maple^ from green^+^ to red^+^ after 15 seconds trans-cuticular exposure to 405 nm light. **C.** Serial imaging of the same M4 NMJ following brief anesthetization at 3 time points (0, 24 hr, 48 hr). PC was performed at timepoint 0 in the 1^st^ instar stage. Although new PSDs incorporated non-synaptic red^+^ GluRIIE during the first day post-PC (middle panel), only non-PC green^+^/red^-^ GluRIIE was present at PSDs that formed after 24 hours (bottom panel). **D.** The percent of red^+^ GluRIIE PSDs remains unchanged between 24 to 48 hours PC, indicating new PSDs formed after 24 hours contain only newly synthesized GluR^New^ (24 hr: 100%, n=15 NMJs in 5 larvae; 48 hr: 101.4% ± 0.85, n=15 in 5 larvae, *p* = 0.11). **E.** Representative single bouton images of PSDs containing GluR^New^ and GluR^Old^ at 24 hours and 144 hours post-PC. **F.** Red^+^ GluR^Old^ fluorescent intensity at larval NMJ PSDs is stable and does not undergo significant decay between 24 to 144 hours post-PC (Day 1: 9070 ± 97.3 average RFU, n=1198 PSDs in 4 larvae; Day 6: 8910 ± 162.2, n=858 PSDs in 4 larvae, *p* = 0.3725). Statistical significance determined with Student’s t-test, ns = not significant. Absolute values and individual statistical tests are summarized in Table S1.

To determine the extent of GluR turnover, average fluorescent intensity of red^+^ GluR^Old^ PSDs was compared at day 1 and day 6 post-PC. Any decay in fluorescence of GluR^Old^ likely reflects turnover, as red-shifted mMaple is highly stable and cannot spontaneously transition back to the green state. No significant decrease in GluR^Old^ fluorescence was observed over this 5-day interval (Fig. 1E, F). These data indicate GluR clusters at *Drosophila* NMJs are highly stable and do not undergo robust turn-over during larval development, consistent with prior imaging of non-mMaple tagged GluRs^15^. The absence of GluR^Old^ at newly formed PSDs later in development indicates GluRs also do not undergo lateral movement between PSDs at *Drosophila* NMJs, similar to observations using mMaple-tagged Cac that is retained within individual presynaptic AZs^16^. Together, these data establish GluRIIE^Maple^ as a tool for time-stamping synapses.

### Synaptic output and release mode are defined by AZ age and maturation state

To examine the effect of age-dependent synapse maturation on presynaptic output, quantal imaging of single SV release events at individual AZs was performed using postsynaptically expressed membrane-tethered myrGCaMP7s as previously described^7,17,18^. Animals expressing myrGCaMP7s and GluRIIE^Maple^ were PC during the early 2^nd^ instar larval stage and quantal imaging was performed four days later to generate AZ *P_r_* maps. The myrGCaMP7s signal is much dimmer than GluRIIE^Maple^ at rest and does not interfere with identification of green^+^ GluR^New^ PSDs. Similarly, the dramatic increase in myrGCaMP7s fluorescence following SV fusion and subsequent postsynaptic opening of GluRs allows simultaneous imaging of synaptic activity on top of baseline GluRIIE^Maple^. The oldest PSDs had more GluR^Old^ and brighter red fluorescence compared to newer PSDs that accumulated only green^+^ GluR^New^ (Fig. 2A). Total GluR^Old^ intensity at individual PSDs was highly correlated with *P_r_* (Pearson r value = 0.56, Fig. 2A, B), indicating evoked output of the corresponding AZ was strongly dependent on synapse age. Comparison of all sites that contained at least some amount of red^+^ GluR^Old^ with newer synapses that lacked any revealed a ∼2.5-fold difference in *P_r_* (Fig. 2C). These data indicate older AZs establish and maintain higher evoked *P_r_* across larval development. However, quantal imaging after four days of PC does not allow precise determination of the output of newer release sites, as GluR^New^ PSDs are indistinguishable whether they formed during day 1 or day 4 after PC. To more precisely examine AZ age and release strength, quantal imaging was performed over a shorter 2-day interval after PC of early 2^nd^ instar larvae (Fig. 2D). A stronger correlation was observed for evoked *P_r_* at sites containing GluR^Old^ versus newly formed PSDs that were less than 1 day old and contained only GluR^New^ (Pearson r value = 0.66, Fig. 2E). Indeed, a >4-fold difference in *P_r_* between older synapses bearing any PC GluRs was observed compared to those without (Fig. 2F).

**Fig. 2.**
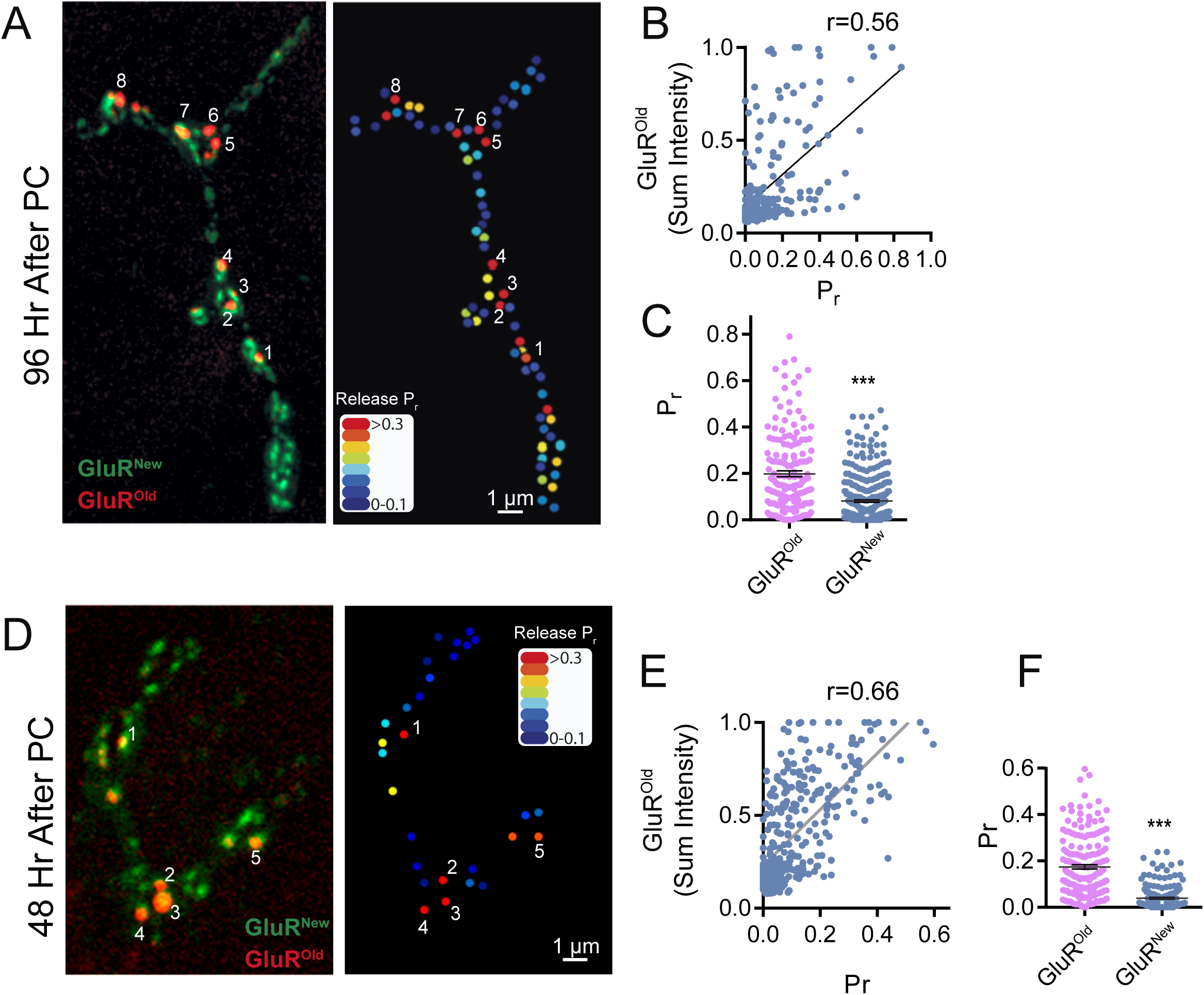
Enhanced evoked synaptic output at older PSDs. **A.** Representative larval M4 NMJ imaged 96 hours after PC with GluR^Old^ PSDs individually numbered (left panel). The corresponding evoked *P_r_* heatmap following quantal imaging with myrGCaMP7s is shown on the right, revealing AZs opposed to the oldest PSDs are among the strongest release sites. **B.** Correlation between PSD age determined by GluR^Old^ fluorescent intensity and evoked *P_r_* (Pearson r value = 0.56). **C.** Average evoked *P_r_* for AZs opposed to older PSDs containing any red^+^ GluR^Old^ compared to younger PSDs with only green^+^/red^-^ GluR^New^ (red^+^ PSDs: 0.193 ± 0.013, n=174 PSDs from 5 larvae; red^-^ PSDs: 0.081 ± 0.005, n=378 PSDs from 5 larvae, *p* < 0.0001). **D.** Representative larval M4 NMJ imaged 48 hours after PC with GluR^Old^ PSDs individually numbered (left panel). The corresponding evoked *P_r_* heatmap following quantal imaging with myrGCaMP7s is shown on the right. **E** Correlation between PSD age determined by red^+^ GluR^Old^ fluorescent intensity and evoked *P_r_* (Pearson r value = 0.66). **F.** Average evoked *P_r_* for AZs opposed to older PSDs containing any red^+^ GluR^Old^ compared to younger PSDs with only green^+^/red^-^ GluR^New^ (red^+^ PSDs: 0.174 ± 0.01, n=179 PSDs from 5 larvae; red^-^ PSDs: 0.0397 ± 0.004, n=192 PSDs from 5 larvae, *p* < 0.0001). Statistical significance determined with Student’s t-test, ns = not significant. Asterisks denote p-values of: ***, p ≤ 0.001. Absolute values and individual statistical tests are summarized in Table S1.

In addition to evoked release, single SVs also fuse through an action-potential independent mechanism to generate spontaneous “mini” events. We previously determined that ∼10% of the AZ population at mature 3^rd^ instar larval NMJs display only spontaneous release and lack evoked fusion^7^. Given the role of maturation for evoked *P_r_*, spontaneous-only AZs might reflect an early developmental stage prior to accumulation of Cac channels required to support evoked release. To determine if spontaneous fusion could occur at AZs lacking Cac, quantification of evoked *P_r_* and spontaneous release rate was performed in larvae expressing endogenously CRISPR-tagged Cac^TagRFP^ ^19^. Indeed, AZs supporting spontaneous SV fusion but lacking evoked release and detectable Cac were present at NMJs (Fig. 3A). Although these data suggest spontaneous-only sites represent an immature state of AZ development, it is unclear if spontaneous release rate is enhanced during AZ maturation as observed for evoked *P_r_*. To correlate synapse age and spontaneous release efficacy, quantal imaging was performed 4 days following PC in GluRIIE^Maple^ larvae. Although a positive correlation between spontaneous release rate and PSD age was observed (Pearson r value = 0.31, Fig. 3B, C), the correlation was significantly less than for evoked *P_r_* (r = 0.56, Fig. 2B). Overall, AZs opposed to sites that contained any red^+^ GluR^Old^ had a ∼2-fold increase in spontaneous release rate compared to sites containing only green^+^ GluR^New^ (Fig. 3D). Similarly, quantifying spontaneous release rate in larvae expressing endogenously CRISPR-tagged Cac^RFP^ revealed that Cac^RFP+^ AZs displayed a 2.2-fold increase in spontaneous release rate over Cac^RFP-^ sites (Fig. 3E). We conclude that spontaneous-only AZs largely represent an early developmental state and that evoked, and to a lesser extent spontaneous release, become more efficient as AZs accumulate scaffolding proteins and VGCCs during maturation.

**Fig. 3.**
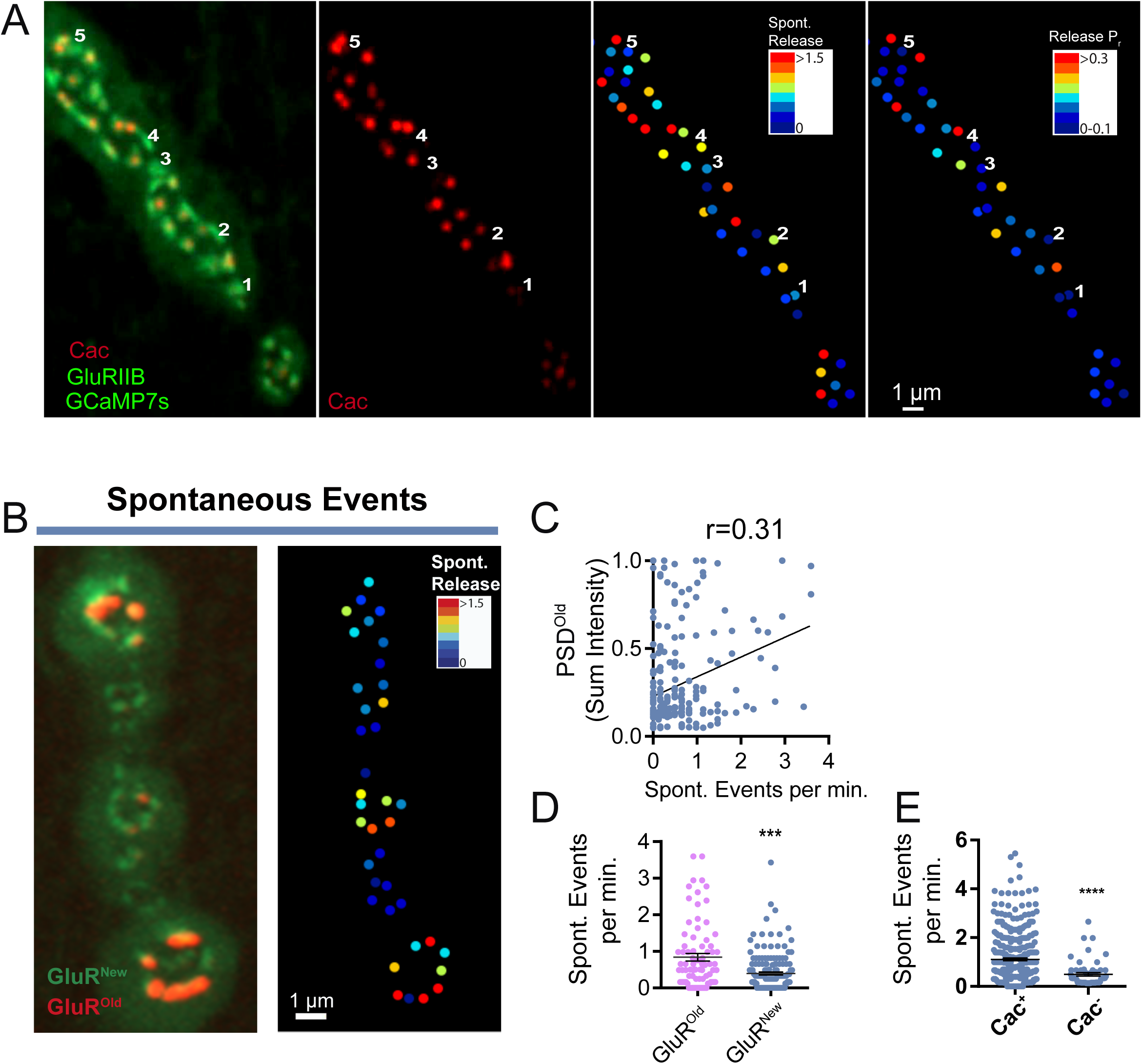
AZs lacking Cac can support spontaneous fusion. **A.** Representative 3^rd^ instar M4 NMJ from a larva expressing GluRIIB-GFP and Cac^RFF^. Several PSDs lacking opposed Cac^RFF+^ signal are individually numbered. Cac^RFF^ signal is shown in panel 2, with spontaneous activity (release events per minute) presented as a heat map at AZs opposed to GluRIIB^+^ PSDs in panel 3. The corresponding evoked *P_r_* heatmap is shown on the right, highlighting lack of evoked release at Cac^-^ AZs. AZs lacking Cac can be functionally silent (sites 2 and 3) or support spontaneous release (sites 1, 4 and 5). **B.** Representative 3^rd^ instar larval M4 NMJ imaged 96 hours after PC to identify GluR^Old^ PSDs (left panel). The corresponding heatmap for spontaneous activity (release events per minute) is shown on the right. **C.** Correlation between PSD age determined by red^+^ GluR^Old^ fluorescent intensity and spontaneous release rate (Pearson r value = 0.31). **D.** Spontaneous release rate for AZs opposed to older PSDs containing any red^+^ GluR compared to younger PSDs with only green^+^/red^-^ GluR (red^+^ synapses: 0.841 ± 0.1 events per min, n=80 AZs from 4 larvae; red^-^ synapses: 0.398 ± 0.04, n=157 AZs from 4 larvae, *p* < 0.0001). **E.** Spontaneous release rate for AZs containing Cac^RFF^ compared to those without (Cac^RFF+^: 1.11 ± 0.05 events per min, n=366 AZs from 5 larvae; Cac^RFF-^: 0.497 ± 0.07, n=55 AZs from 5 larvae, *p* < 0.0001). Statistical significance determined with Student’s t-test, ns = not significant. Asterisks denote p-values of: ***, p ≤ 0.001 and ****, p ≤ 0.0001. Absolute values and individual statistical tests are summarized in Table S1.

### Effects of NMJ growth and *rab3* mutants on AZ maturation and output

During larval growth, NMJs undergo significant expansion, with addition of new AZs and co-opposed PSDs. Prior trans-cuticular imaging during early larval stages revealed an approximate doubling of AZ number each day (∼1.7-fold increase)^7^. To precisely quantify synapse addition across larval development, the ratio of GluR^New^ versus GluR^Old^ PSDs was determined at different developmental points. When PC was performed in the early 1^st^ instar and synapse addition was measured 24 to 48 hours later, a large increase in synapse number was observed (1.78-fold, Supplemental Fig. 1A). When PC was performed later in the 3^rd^ instar period and synapse addition was measured between day 5 to 6 of development, the rate of synapse addition had slowed (1.33-fold). These data indicate NMJ growth and AZ addition is nonuniform across development and decreases during the later 3^rd^ instar stage.

Prior studies revealed a heterogeneity in synaptic transmission strength along some *Drosophila* larval motor axons, with terminal boutons showing enhanced Ca^2+^ influx and release^20^. This graded transmission was not uniform across all MNs, as M6/7 MNs displayed enhanced output at terminal boutons while M4 MNs lacked this effect^21^. Molecular mechanisms explaining enhanced release from terminal boutons and differences across MN types remain unclear. Given NMJ expansion can result in individual release sites changing their relative position during larval development (Supplemental Fig. 1B), we hypothesized distinct patterns of bouton addition and synaptic growth might contribute to the observed release heterogeneity. To compare patterns of NMJ growth at M6/7 versus M4, the relative abundance of GluR^Old^ PSDs was quantified at individual boutons from proximal to distal after several days of growth post-PC (Supplemental Fig. 1C, D). Axon terminals of M6/7 MNs commonly grew by “stretching” and adding new internal boutons such that terminal boutons contained more GluR^Old^ PSDs. In contrast, axons innervating M4 often grew by budding new branches that resulted in new terminal boutons lacking or having fewer older PSDs. The distinct pattern of synaptic growth and ∼2.5-fold enhancement of mature synapses at terminal boutons of M6/7 NMJs provides an underlying mechanism to explain differences in graded transmission between these two MN subtypes.

The correlation between synapse age and evoked release suggests *P_r_* heterogeneity reflects age-dependent maturation of release output. Mutations in *Drosophila rab3* cause a striking defect in AZ material accumulation, with Cac channels and late scaffolds like BRP hyperaccumulating at a subset of AZs, while other AZs are devoid of these components^22^. One hypothesis for the *rab3* phenotype is that the earliest forming AZs require a distinct maturation mechanism coupled to axonal pathfinding and target recognition that is Rab3-independent. Addition of late scaffolds and Cac to AZs formed during subsequent NMJ expansion would then require Rab3 function. This model predicts only AZs formed in the embryo and 1^st^ instar stage would contain late AZ material in *rab3* mutants, and that evoked *P_r_* heterogeneity would be reduced given mature AZs would be more age-matched than in controls. To test this hypothesis, GluRIIE^Maple^ was expressed in *rab3* mutants and PC was performed at different larval stages. Larvae were subsequently immunostained to determine the presence of the late scaffold BRP at 3^rd^ instar AZs. If BRP^+^ AZs were only opposed to GluR^Old^ PSDs, it would argue that BRP cannot accumulate at AZs formed later in development. Consistent with this model, BRP co-localized with just 8.5% of opposed PSDs that only contained GluR^New^ in *rab3* mutants when PC was performed at the 1^st^ instar stage compared to 87% in controls (Supplemental Fig. 1E, F). Larvae PC at the 2^nd^ instar stage to ensure labeling of only the later wave of new synapses revealed an even lower percent (0.9%) of BRP^+^ AZs opposed to GluR^New^ PSDs in *rab3* mutants. Consistent with AZ age and material accumulation correlating with evoked *P_r_*, quantal imaging in *rab3* mutants demonstrated *P_r_* distribution was more homogenous and shifted towards higher *P_r_* than controls (Supplemental Fig. 1G). These data indicate early formed AZs contain late scaffolds and Cac channels in *rab3* mutants, with AZs formed later in development requiring Rab3 to accumulate these proteins.

### Sequential addition and accumulation of AZ proteins at nascent synapses

Although a number of transport mechanisms are implicated in material delivery to synapses^23–25^, it is unclear if most AZ components are co-trafficked and delivered as pre-assembled modules. At *Drosophila* NMJs, a sequential process of AZ assembly has been suggested, with Liprin-α and Syd1 arriving early during AZ formation^26^. To quantify the pattern of accumulation for a broad range of AZ proteins, fluorescently-tagged lines expressing Liprin-α, Syd1, BRP, RIM, RBP, Cac, Unc13B and Unc13A were analyzed at early 2^nd^ instar larval NMJs when AZ addition is robust. The presence of these proteins across the AZ population was correlated with the distribution of endogenously CRISPR-tagged Cac channels (Cac^GFP^ or Cac^RFP^) and endogenous RBP, a late AZ scaffold, detected by antibody staining (Fig. 4A). Antisera to RBP served as a reliable marker for endogenous RBP and co-stained 99.3% of RBP^+^ AZs expressing exogenous fluorescently-tagged RBP. Liprin-α was the earliest arriving AZ protein with 34.1% of Liprin-α^+^ puncta lacking RBP and 43.4% lacking Cac. To determine if these Liprin-α^+^ puncta were newly formed AZs or accumulations of the protein outside of release sites, the presence of co-opposed postsynaptic GluRIIB^+^ receptor fields was quantified. 93.3% of Liprin-α puncta were opposed to GluRIIB^+^ PSDs (Fig. 4B), indicating most Liprin-α^+^ clusters represent AZs in various states of maturation. RIM and Syd1 also arrived early to AZs, with 9.7% of RIM^+^ AZs lacking RBP and 24.3% lacking Cac, while 9.6% of Syd1^+^ AZs lacked RBP and 18.1% lacked Cac (Fig. 4A). In contrast, BRP arrived later, with only 1.6% of BRP^+^ AZs lacking RBP and 11% lacking Cac. To assay the arrival and distribution of the two Unc13 spicing isoforms^6^, CRISPR was used to endogenously tag Unc13B with mClover and Unc13A with mRuby. Unc13B arrived early to AZs as previously observed^6^, with 10.8% of Unc13B^+^ AZs lacking RBP and 19.6% lacking Cac. Unc13A followed a similar pattern to BRP, with only 1.7% of Unc13A^+^ AZs lacking RBP and 9% lacking Cac (Fig. 4A).

**Fig. 4.**
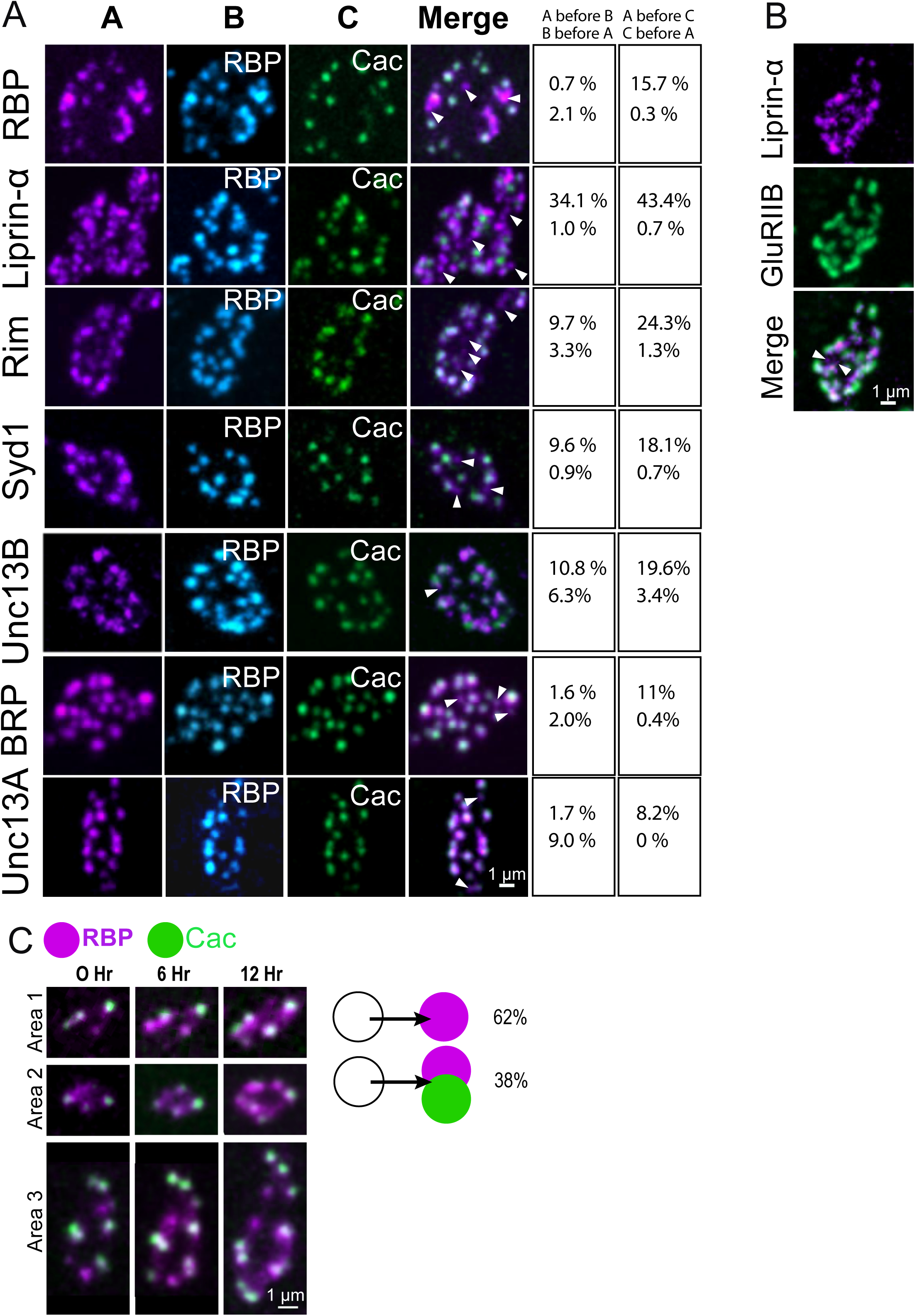
Time course for protein accumulation during AZ maturation. **A.** Representative 2^nd^ instar M4 synaptic boutons with an analysis of the distribution of fluorescent puncta for the indicated AZ proteins (left panel A) in relation to immunostained RBP (middle panel B) and endogenous CRISPR GFP-tagged or RFP-tagged Cac (right panel C). The merged image shows labeling for Cac and the AZ protein indicated in panel A, with representative AZs lacking Cac indicated by white arrows. The right panels show the % of individual puncta for each AZ protein lacking RBP or Cac (top values) versus RBP or Cac puncta lacking the indicated AZ protein (bottom values). N=7 NMJs from 2 larvae for each correlation. **B.** Representative image of a 2^nd^ instar M4 synaptic bouton showing Liprin-α puncta compared to the distribution of GluRIIB. Most puncta overlap, although a few are positive for Liprin-α only (white arrows). **C.** Serial imaging of RBP and Cac appearance at 2^nd^ instar M4 AZs over 3 time points indicate RBP accumulates before Cac arrival. The appearance of new RBP or Cac accumulations within 6-hour intervals was scored (n=10 NMJs from 3 larva), revealing 62% of newly formed puncta are only RBP^+^ and 38% are RBP^+^ and Cac^+^. No case of Cac at newly formed AZs lacking RBP was observed. Absolute values are summarized in Table S1.

Although differential accumulation of proteins observed in fixed imaging suggests a sequential AZ maturation process, it doesn’t provide a time-course for AZ assembly. To examine AZ maturation over time, serial imaging of briefly anesthetized larvae expressing fluorescently-tagged RBP and Cac was performed at 0, 6 and 12 hour time points (Fig. 4C). The appearance of new RBP^+^ AZ clusters and their subsequent accumulation of Cac was scored across the three imaging sessions. For 62% of cases where new RBP clusters appeared during a 6-hour imaging window, Cac was not detected at these AZs. Newly formed RBP^+^ clusters that also contained Cac appeared 38% of the time, indicating ∼4 hours is required for Cac channels to arrive after RBP accumulation. Together, these data suggest a sequential addition of AZ proteins whereby Liprin-α arrives first, followed by a cohort of early scaffolds (Syd1, RIM, and Unc13B). BRP, RBP and Unc13A arrive at AZs later and around the same time, with the VDCC Cac following several hours after late scaffolds accumulate.

### Early AZ scaffold levels correlate poorly with evoked release, while late AZ scaffold abundance predicts presynaptic strength

Given sequential addition of proteins to maturing AZs, how their abundance correlates with synapse function was assayed. Presynaptic release strength has been previously correlated with AZ abundance of Ca^2+^ channels^7,8,19^, while late AZ scaffolding proteins like RBP and BRP facilitate clustering of Cac at *Drosophila* AZs^11,27,28^. Whether the abundance of early synaptic scaffolds also regulates release output is unclear. To examine early versus late scaffold abundance and presynaptic output, fluorescently-tagged lines expressing Liprin-α, Syd1, RBP and Unc13A were assayed for Cac levels and evoked *P_r_*. AZ Cac levels correlated weakly with the abundance of the early scaffolding proteins Liprin-α (Pearson r value = 0.19) and Syd1 (Pearson r value = 0.26, Fig. 5A-C). In contrast, AZ abundance of the late scaffolding proteins RBP (Pearson r value = 0.51) and Unc13A (Pearson r value = 0.66) were more strongly correlated with Cac (Fig. 5A, D, E). To assay AZ protein abundance and evoked release strength, functional imaging was performed in larvae expressing LexOP-myrGCaMP7s in muscles with *Mef2*-LexA and a UAS-fluorescently-tagged AZ protein in MNs using *elav*-GAL4. As suggested by their low correlation with Cac abundance, AZ levels of Liprin-α (Pearson r value = 0.13, Fig. 5F) and Syd1 (Pearson r value = 0.12, Fig. 5G) poorly predicted evoked *P_r_* compared to the AZ abundance of RBP (Pearson r value = 0.37, Fig. 5H) and Unc13A (Pearson r value = 0.52, Fig. 5I). Although late scaffolds correlated with evoked output, their abundance was slightly less predictive of *P_r_* that Cac AZ levels (Pearson r value = 0.54, Fig. 5J). These data indicate that although early AZ scaffolds regulate AZ seeding, their abundance at mature release sites is less critical for Cac levels and *P_r_* than late-arriving AZ proteins like RBP and Unc13A.

**Fig. 5.**
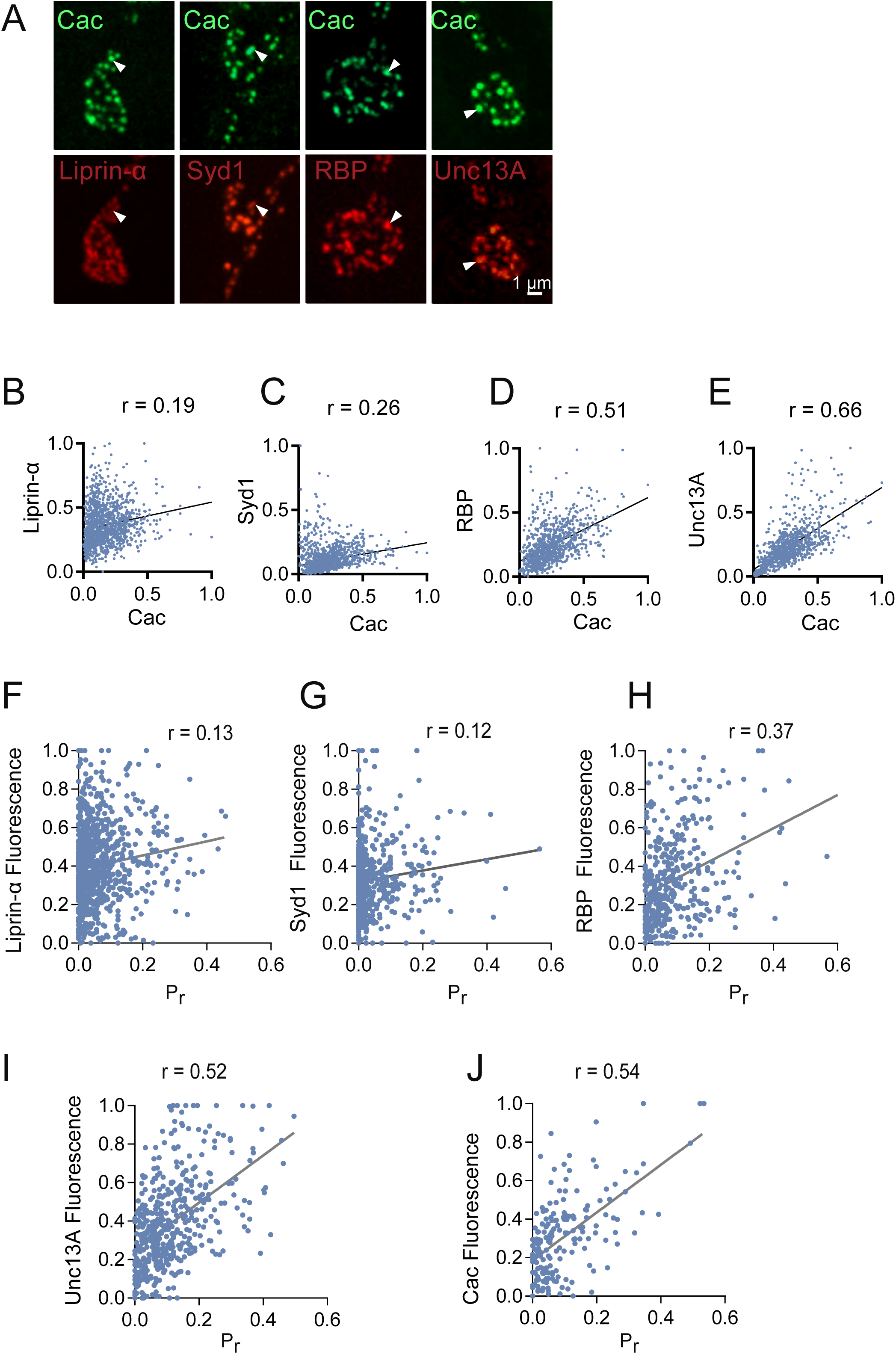
The abundance of early AZ scaffolds poorly predicts Cac accumulation and *P_r_*. **A.** Representative images of 3^rd^ instar M4 synaptic boutons expressing endogenous CRISPR-tagged Cac^GFP^ (top row) and the indicated AZ scaffolding protein (bottom row). One of the brightest Cac^+^ puncta in each bouton is highlighted (white arrows) to compare abundance to the indicated scaffolding protein. **B-E.** Correlation between AZ abundance (fluorescent intensity) of Cac and Liprin-α (**B**, n = 1422 AZs from 6 NMJs from 6 larvae), Syd1 (**C**, n = 1579 AZs from 6 NMJs from 6 larvae), RBP (**D**, n = 996 AZs from 6 NMJs from 6 larvae) and Unc13A (**E**, n = 984 AZs from 6 NMJs from 6 larvae) at 3^rd^ instar M4 NMJs. **F-J.** Correlation between evoked *P_r_* and AZ abundance (fluorescent intensity) of Liprin-α **(F**, n = 913 AZs from 6 NMJs from 6 larvae), Syd 1 **(G**, n = 525 AZs from 6 NMJs from 6 larvae), RBP **(H**, n = 407 AZs from 6 NMJs from 6 larvae), Unc13A **(I**, n = 462 AZs from 8 NMJs from 6 larvae) and Cac **(J**, n = 486 AZs from 7 NMJs from 6 larvae) at 3^rd^ instar M4 NMJs. Absolute values and individual statistical tests are summarized in Table S1.

### Changes in neuronal excitability regulate AZ and PSD size

Alterations in synaptic output occur through both Hebbian and homeostatic mechanisms in response to changes in neuronal activity. At *Drosophila* NMJs, the segregation of GluRs within a PSD (stronger GluRIIA^+^-containing receptors at the center versus weaker GluRIIB^+^ at the periphery) is enhanced during early stages of synapse formation following elevated neuronal activity^7^. How changes in neuronal activity influence presynaptic AZ maturation and structure is less clear, though recent studies suggest AZ remodeling occurs during homeostatic plasticity^29^. To assay changes in AZ formation in response to altered neuronal activity, AZ protein area and accumulation were examined following manipulations that decrease or increase synaptic output. Synaptotagmin 1 (Syt1) functions as the Ca^2+^ sensor for synchronous SV fusion and *syt1^-/-^* null mutants display a severe reduction in evoked release^30^. Analysis of the distribution of late AZ scaffolds and Cac at 3^rd^ instar larval M4 NMJs in *syt1^-/-^* revealed a ∼15% enlargement in AZ Cac area compared to controls (Fig. 6A, B). Changes in Cac area in *syt1^-/-^* were accompanied by a mild reduction in mean AZ Cac brightness, although total Cac AZ abundance was not different (Fig. 6C, D). To examine the effects of increased neuronal activity, double mutants lacking the Shaker (Sh) and Eag delayed rectifier K^+^ channels that enhance release^31^ were analyzed. In contrast to enlarged AZs in *syt1^-/-^*, *Sh^KS133^ eag*^1^ mutants showed a ∼11% reduction in AZ Cac area (Fig. 6A, B). The mean AZ brightness for both Cac and BRP was increased without affecting the total amount of those proteins in *Sh^KS133^ eag^1^* (Fig. 6C, D), suggesting AZ components are distributed over a larger or smaller AZ area in response to altered neuronal activity. Together, these data indicate AZ size and protein clustering is sensitive to changes in neuronal activity, as observed for AZ protein density changes during homeostatic plasticity following disruptions to postsynaptic GluR function^4,32,33^.

**Fig. 6.**
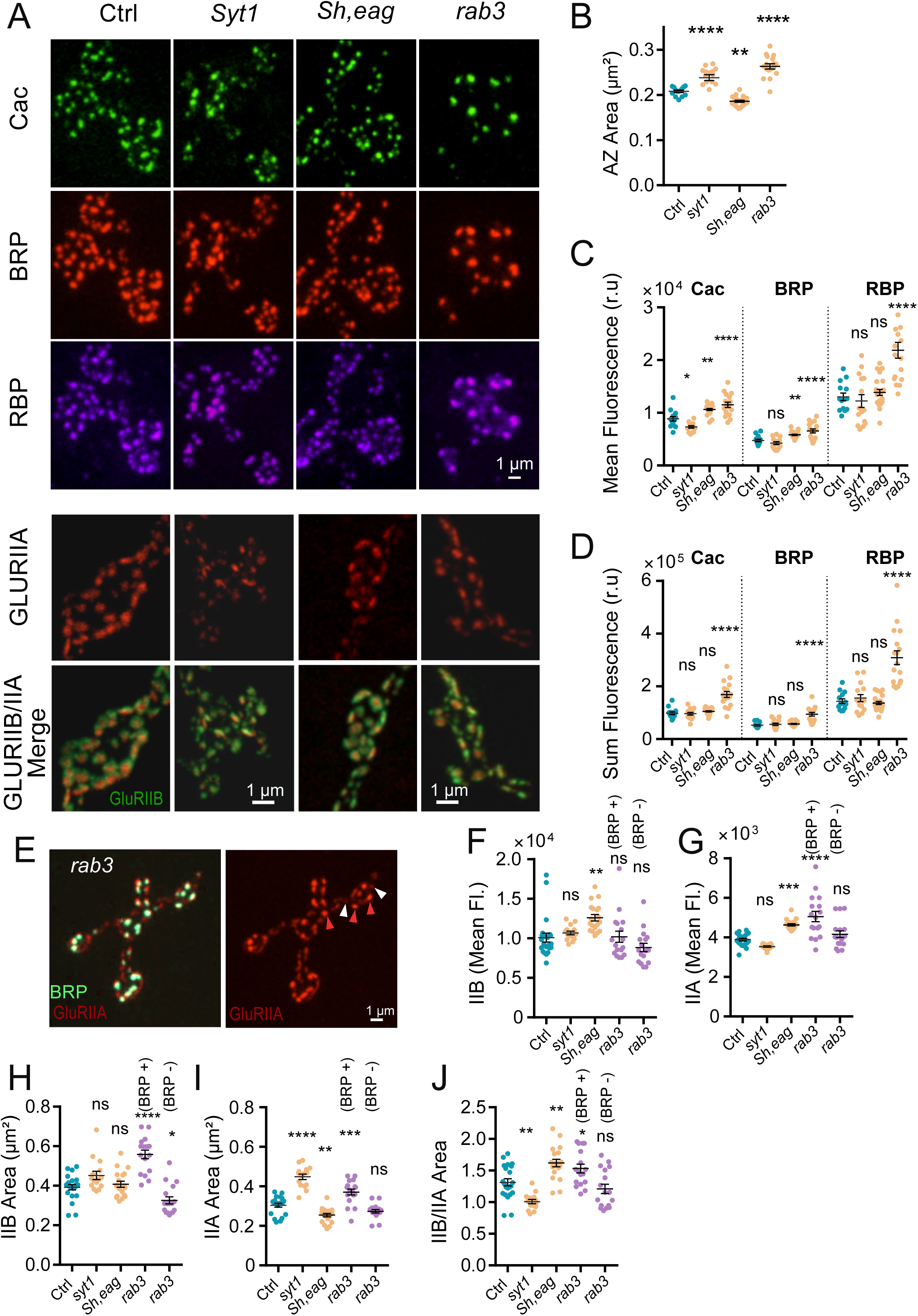
Increased or decreased synaptic output alter AZ area and AZ/PSD material distribution. **A.** Representative images of presynaptic (Cac, BRP, RBP) and postsynaptic (GluRIIA and GluRIIB) proteins at 3^rd^ instar M4 NMJs in control, *syt1^AD4/N13^*, *Sh^KS133^ eag^1^*, and *rab3^rup^* mutants. **B.** Analysis of AZ Cac area for the indicated genotypes (Ctrl: n=14 NMJs from 4 larvae*; syt1^-/-^*: n=15 NMJs from 4 larvae, *p* < 0.0001; *Sh^KS133^ eag^1^*: n=23 NMJs from 6 larvae, *p* = 0.0022; *rab3^rup^*: n=17 NMJs from 5 larvae, *p* < 0.0001). **C.** Mean fluorescent intensity per AZ for Cac, BRP and RBP for the indicated genotypes (Ctrl: n=14 NMJs from 4 larvae*; syt1^-/-^*: n=15 NMJs from 4 larvae; *Sh^KS133^ eag^1^*: n=23 NMJs from 6 larvae; *rab3^rup^*: n=17 NMJs from 5 larvae; *syt1^-/-^* Cac: *p* = 0.0144; *Sh^KS133^ eag^1^* Cac: *p* = 0.0014; *rab3^rup^* Cac: *p* < 0.0001; *syt1^-/-^* BRP: *p* = 0.429; *Sh^KS133^ eag^1^* BRP: *p* = 0.0115; *rab3^rup^* BRP: *p* < 0.0001; *syt1^-/-^* RBP: *p* = 0.924; *Sh^KS133^ eag^1^* RBP: *p* = 0.888; *rab3^rup^* RBP: *p* < 0.0001). **D.** Sum fluorescent intensity per AZ for Cac, BRP and RBP for the indicated genotypes (Ctrl: n=14 NMJs from 4 larvae*; syt1^-/-^*: n=15 NMJs from 4 larvae; *Sh^KS133^ eag^1^*: n=23 NMJs from 6 larvae; *rab3^rup^*: n=17 NMJs from 5 larvae; *syt1^-/-^* Cac: *p* = 0.9952; *Sh^KS133^ eag^1^* Cac: *p* = 0.7896; *rab3^rup^* Cac: *p* < 0.0001; *syt1^-/-^* BRP: *p* = 0.9; *Sh^KS133^ eag^1^* BRP: *p* = 0.706; *rab3^rup^* BRP: *p* < 0.0001; *syt1^-/-^* RBP: *p* = 0.918; *Sh^KS133^ eag^1^* RBP: *p* = 0.978; *rab3^rup^* RBP: *p* < 0.0001). **E.** Representative 3^rd^ instar M4 NMJ in *rab3^rup^* immunostained for BRP and GluRIIA showing differential accumulation of GluRIIA at PSDs opposed to presynaptic AZs with or without BRP (white arrows). **F.** Mean fluorescent intensity of GluRIIB for the indicated genotypes, with BRP^+^ versus BRP^-^ AZs separately quantified for *rab3* mutants (Ctrl: n=22 NMJs from 6 larvae*; syt1^-/-^*: n=15 NMJs from 4 larvae, *p* = 0.848; *Sh^KS133^ eag^1^*: n=19 NMJs from 5 larvae, *p* = 0.0021; *rab3^rup^* n=17 NMJs from 5 larvae, BRP^+^ *p* = 0.999, BRP^-^ *p* = 0.2587). **G**. Mean fluorescent intensity of GluRIIA for the indicated genotypes (Ctrl: n=22 NMJs from 6 larvae*; syt1^-/-^*: n=15 NMJs from 4 larvae, *p* = 0.242; *Sh^KS133^ eag^1^*: n=19 NMJs from 5 larvae, *p* = 0.0005; *rab3^rup^* n=17 NMJs from 5 larvae, BRP^+^ *p* < 0.0001, BRP^-^ *p* = 0.414). **H.** Mean GluRIIB area for the indicated genotypes (Ctrl: n=22 NMJs from 6 larvae*; syt1^-/-^*: n=15 NMJs from 4 larvae, *p* = 0.064; *Sh^KS133^ eag^1^*: n=19 NMJs from 5 larvae, *p* = 0.912; *rab3^rup^* n=17 NMJs from 5 larvae, BRP^+^ *p* < 0.0001, BRP^-^ *p* = 0.025). **I.** Mean GluRIIA area for the indicated genotypes (Ctrl: n=22 NMJs from 6 larvae*; syt1^-/-^*: n=15 NMJs from 4 larvae, *p* < 0.0001; *Sh^KS133^ eag^1^*: n=19 NMJs from 5 larvae, *p* = 0.005; *rab3^rup^* n=17 NMJs from 5 larvae, BRP^+^ *p* = 0.0001, BRP^-^ *p* = 0.181). **J.** Ratio of GluRIIB/GluRIIA area for the indicated genotypes (Ctrl: n=22 NMJs from 6 larvae*; syt1^-/-^*: n=15 NMJs from 4 larvae, *p* = 0.0026; *Sh^KS133^ eag^1^*: n=19 NMJs from 5 larvae, *p* = 0.0011; *rab3^rup^* n=17 NMJs from 5 larvae, BRP^+^ *p* = 0.035, BRP^-^ *p* = 0.537). Statistical significance determined with one-way ANOVA followed by Tukey’s multiple comparisons test. Asterisks denote p-values of: *, p ≤ 0.05; **, p ≤ 0.01; ***, p ≤ 0.001; and ****, p ≤ 0.0001. Absolute values and individual statistical tests are summarized in Table S1.

We next examined how activity-dependent changes in AZ structure compared to those in *rab3* mutants. Loss of Rab3 results in accumulation of late scaffolding proteins and Cac at a subset of AZs^22^, substantially increasing AZ area at these sites (Fig. 6A). The enlargement of AZ size in *rab3* mutants is not likely due to enhanced release since *Sh^KS133^ eag^1^* mutants displayed reduced AZ area. Enlarged AZs in *rab3* may instead reflect distribution of proteins across a smaller number of release sites^34^. Although *rab3* mutants do not provide an ideal system to assay activity-dependent presynaptic plasticity due to the protein’s role in AZ material distribution, it is a valuable tool to characterize PSD changes within the same postsynaptic cell for sites lacking evoked release (BRP^-^/Cac^-^) versus those with higher *P_r_* (BRP^+^/Cac^+^). Prior data indicated spatial segregation of GluRIIA and GluRIIB within single PSDs was enhanced in newly forming synapses experiencing elevated activity, with GluRIIA enriching at the PSD center and GluRIIB forming a peripheral ring around GluRIIA^7^. As these experiments were performed in live larvae, the intensity of GluRIIA and GluRIIB could not be directly compared over imaging sessions due to changes in cuticle thickness across development. To examine how GluR levels are affected by the activity of their opposed AZ, GluRIIA intensity in *rab3* mutants was quantified at PSDs opposed to Cac^-^ or Cac^+^ release sites in *rab3* mutants. A 21% increase in average GluRIIA intensity was observed at PSDs opposed to Cac^+^ AZs (Fig. 6E, G), similar to prior observations showing enhanced GluRIIC levels at BRP^+^ AZs in *rab3* mutants^22^. *Syt1^-/-^* mutants did not show a significant decrease in GluRIIA levels, while *Sh^KS133^ eag^1^* mutants displayed elevated GluRIIA similar to *rab3* BRP^+^/Cac^+^ release sites (Fig. 6A, G). These data indicate enhanced release is the primary driver for increased GluRIIA abundance at Cac^+^ AZs in *rab3* mutants.

Although GluRIIB intensity was not significantly different across AZs in *rab3* mutants (Fig. 6F), GluRIIB PSD area was increased at Cac^+^ AZs (Fig. 6H). Indeed, AZs with elevated activity (*rab3* Cac^+^ and *Sh^KS133^ eag^1^*) had the highest ratio of GluRIIB to GluRIIA area (Fig. 6I, J), indicating elevated activity enhances GluR segregation to concentrate GluRIIA in the PSD center and GluRIIB at the periphery. A smaller GluRIIB/GluRIIA area was observed at AZs with reduced activity (*rab3* Cac^-^ and *syt1^-/-^*) compared to controls (Fig. 6J). Together, these data indicate neuronal activity alters the size of both presynaptic AZs and postsynaptic GluR fields, while also regulating GluRIIA levels and segregation within the PSD. We hypothesize elevated activity activates a positive feedback loop during early stages of synapse formation to enhance accumulation of synaptic components. At later stages when *P_r_* is established, a negative feedback signal arrests further AZ enlargement and material accumulation. When synaptic activity is reduced as observed in *syt1^-/-^* mutants, negative feedback is suppressed and AZs continue to accumulate material and increase in size.

### PSD GluRIIA accumulation is enhanced by activity early in synapse development

To determine which stage of synaptic development is most sensitive to neuronal activity, GluRIIA intensity was compared at PSDs opposed to Cac^-^ versus Cac^+^ release sites in *rab3* mutants across larval development. The difference in average GluRIIA intensity at PSDs opposed to functional (Cac^+^) versus nonfunctional (Cac^-^) AZs was strongest in the 2^nd^ instar (1.98-fold) and declined at the late 3^rd^ instar stage (1.2-fold, Fig. 7A, B), suggesting a more prominent role for activity earlier in development. To examine synaptic development independent of *rab3* mutants, Cas9 and UAS-*Cac* guide RNAs were expressed in MN1-Ib that innervates M1 using a MN1-Ib-Gal4 driver^18^. This approach bypasses confounding effects of whole-animal activity manipulations by restricting synaptic dysfunction to a single MN in each hemi-segment of the larvae. This single neuron CRISPR approach resulted in disruption of Cac expression in MN1-Ib late in embryogenesis, with a subset of early-formed AZs containing residual Cac, and AZs formed later in development lacking the protein (Fig. 7C). Overall, only 34.4% of AZs opposed to GluRIIA^+^ PSDs contained residual Cac at M1 NMJs, compared to 88.3% of AZs at unaffected M9 NMJs. The distribution of synaptic proteins at different larval stages was compared at M1 NMJs with reduced activity to unaffected NMJs on neighboring M9 in the same animal. At the 2^nd^ instar stage, GluRIIA abundance was decreased at PSDs of M1 NMJs versus those of M9 (Fig. 7D), consistent with synaptic activity promoting GluRIIA PSD accumulation. Although the strong correlation between Cac and GluRIIA abundance at individual release sites observed at M9 NMJs was reduced at M1, AZs that contained residual Cac still displayed higher GluRIIA PSD levels than Cac^-^ AZs (Fig. 7E). In addition to reduced levels, GluRIIA also failed to concentrate in the PSD center at M1 NMJs with reduced activity, leading to a larger GluRIIA PSD area (Fig. 7F) as observed in *syt1^-/-^* mutants (Fig. 6I). These data indicate the rate of segregation of GluRIIA to the PSD center is reduced at inactive synapses. Although the enlarged GluRIIA PSD area persisted into the 3^rd^ instar stage (Fig. 7I), PSDs at M1 NMJs accumulated GluRIIA to similar levels as M9 NMJs at this later developmental stage (Fig. 7J). Overall, the postsynaptic side displayed larger area changes to activity reduction at Cac^-^ AZs, as BRP area was less dramatically enlarged compared to GluRIIA fields (Figure 7K). We conclude that GluRIIA accumulation and segregation within the PSD is activity-dependent early in synapse formation, with silent PSDs continuing to accumulate GluRIIA over development at a slower rate. Together with differences between Cac^+^ versus Cac^-^ AZs in *rab3* mutants, these data indicate enhanced GluRIIA PSD accumulation and segregation early in synapse formation reflects the output of the opposed AZ rather than integrated activity across the entire NMJ.

**Fig. 7.**
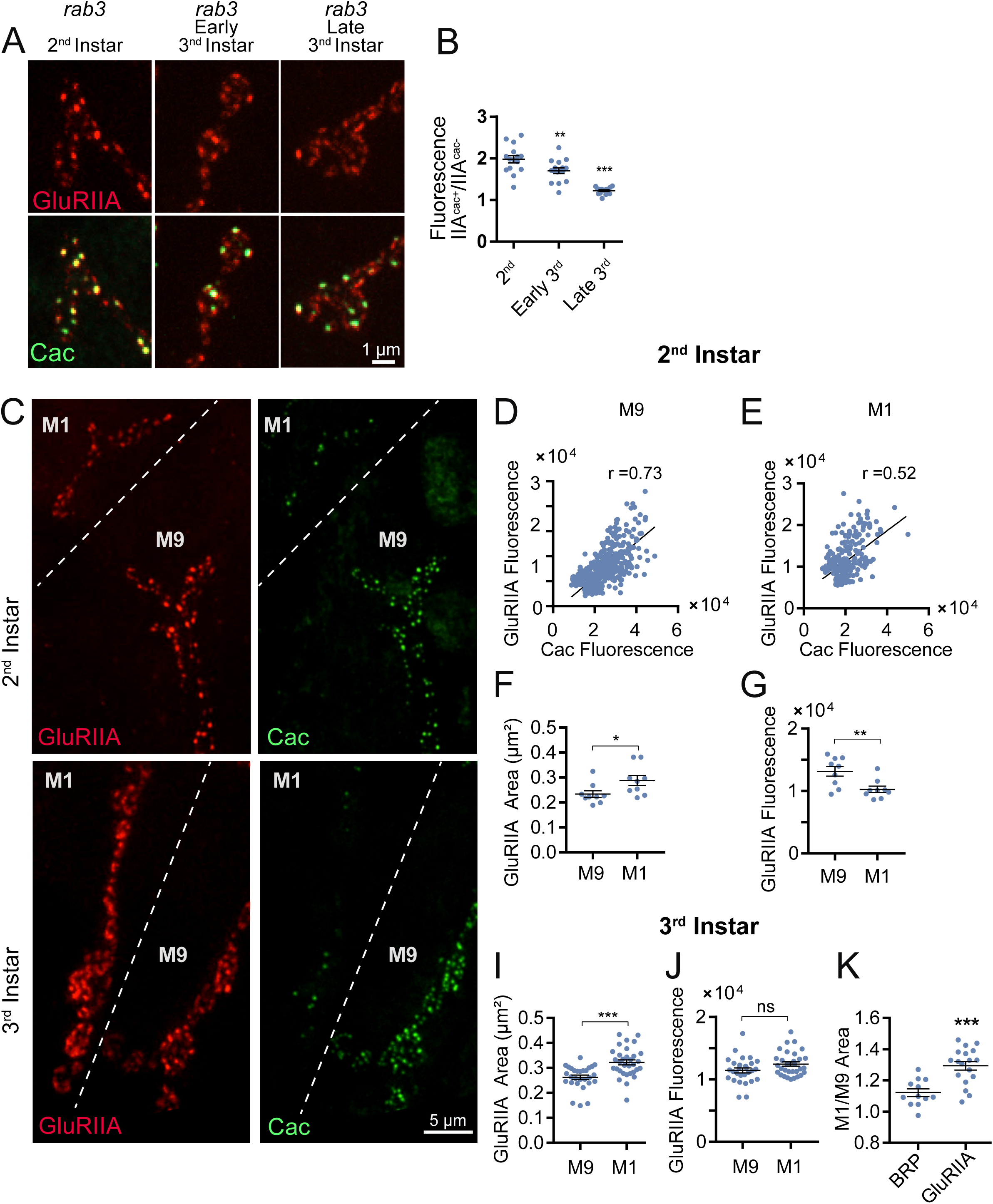
GluRIIA abundance and segregation is sensitive to altered output in *Rab3* mutants and following CRISPR-mediated *Cac* deletion. A. Representative images of GluRIIA and Cac immunolabelling within synaptic boutons at M4 NMJs of *rab3* mutants at three developmental stages. **B.** Quantification of the ratio of GluRIIA fluorescent intensity at PSDs opposed to Cac^+^ versus Cac^-^ AZs across 3 developmental stages (2^nd^ instar: n=15 NMJs from 4 larvae; early 3^rd^ instar: n=16 NMJs from 4 larvae, *p* = 0.0073; late 3^rd^ instar: n=15 NMJs from 4 larvae, *p* < 0.0001). **C.** Representative images of GluRIIA and Cac immunolabelling at 2^nd^ (top panel) and 3^rd^ (bottom panel) instar stages for reduced output M1 NMJs versus unaffected M9 NMJs in MN1-Ib-Gal4; UAS-Cas9, UAS-*Cac* gRNA expressing larvae. **D, E.** Correlation between AZ Cac and PSD GluRIIA fluorescence in 2^nd^ instar MN1-Ib-Gal4; UAS-Cas9, UAS-*Cac* gRNA expressing larvae for M9 (**D**, n=616 AZ/PSDs from 15 NMJs from 4 larvae, Pearson r value = 0.73) and M1 NMJs (**E**, n=299 AZ/PSDs from 15 NMJs from 4 larvae, Pearson r value = 0.52). **F, G.** Quantification of GluRIIA PSD area **(F**, *p* = 0.039) and mean GluRIIA fluorescence (**G**, *p* = 0.007) in 2^nd^ instar MN1-Ib-Gal4; UAS-Cas9, UAS-*Cac* gRNA expressing larvae for M9 and M1 NMJs (n=9 NMJs from 3 larvae). **I, J.** Quantification of GluRIIA PSD area **(I**, *p* < 0.0001) and mean GluRIIA fluorescence (**J**, *p* = 0.054) in 3^rd^ instar MN1-Ib-Gal4; UAS-Cas9, UAS-*Cac* gRNA expressing larvae for M9 and M1 NMJs (n=28 NMJs from 4 larvae). **K.** Quantification of the M1/M9 ratio of AZ BRP area and PSD GluRIIA area in 3^rd^ instar MN1-Ib-Gal4; UAS-Cas9, UAS-*cac* gRNA expressing larvae (BRP: n=12 NMJs from 3 larvae; M1: n=18 NMJs from 4 larvae, *p* = 0.0001). Statistical significance determined with one-way ANOVA followed by Tukey’s multiple comparisons test. Asterisks denote p-values of: ***, p ≤ 0.001. Absolute values and individual statistical tests are summarized in Table S1.

### Reduced synaptic activity increases AZ size via a Rab3-independent mechanism

To further characterize changes in AZ development following disruption to synaptic transmission, single MN manipulations of synaptic output were performed using the MN1-Ib Gal4 driver. Pan-neuronal expression of a dominant-negative *Syt1* transgene that blocks Ca^2+^ binding by the C2B domain (UAS-Syt1C2B^D416N,D418N^, hereafter as Syt1^DN^) causes lethality and reduces evoked release to a greater extent than observed in *syt1^-/-^* null mutants^35^. Similarly, pan-neuronal expression of tetanus toxin light chain (hereafter referred to as UAS-TeNT) cleaves the v-SNARE n-Synaptobrevin (nSyb), causing lethality and abolishing evoked release^36^. TeNT or Syt1^DN^ were expressed in MN1-Ib and the distribution of BRP, RBP and Cac was analyzed and compared to control NMJs. AZ Cac area was increased by 47% following TeNT expression and 21% following Syt1^DN^ expression (Fig. 8A, B). In contrast to *syt1^-/-^* mutants where more evoked release persists, AZ accumulation of BRP, RBP and Cac was also enhanced at MN1-Ib NMJs expressing either TeNT or Syt1^DN^ (Figure 8C-E).

**Fig. 8.**
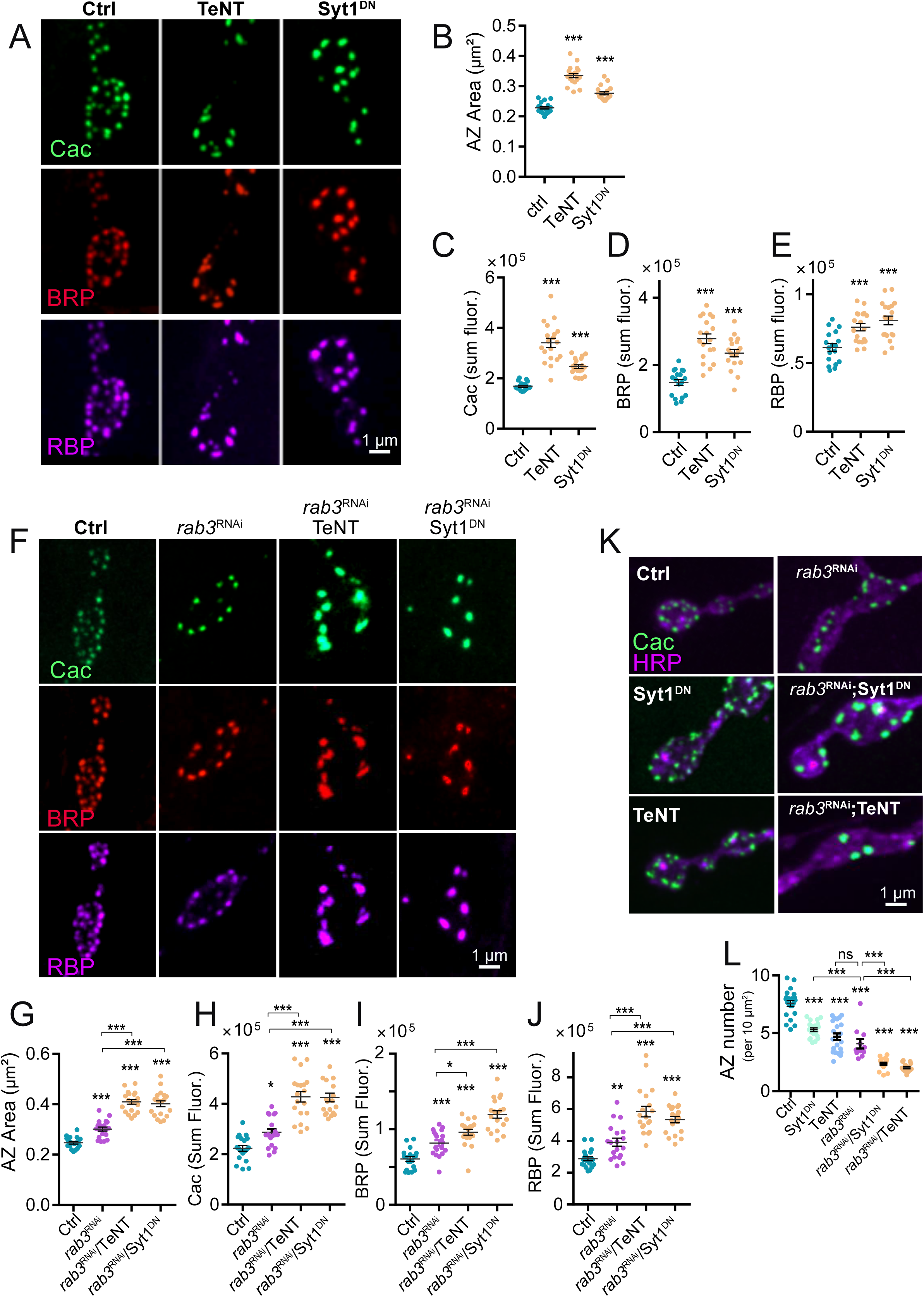
Presynaptic silencing results in AZ enlargement via a Rab3-independent mechanism. **A.** Representative images of Cac, BRP and RBP immunolabelling within synaptic boutons of 3^rd^ instar M1 NMJs of controls or following expression of TeNT (middle panel) or Syt1^DN^ (right panel) with the MN1-Ib-Gal4 driver. **B**. Quantification of AZ Cac area in the three genotypes (Ctrl: n=18 NMJs from 5 larvae; TeNT: n=19 NMJs from 5 larvae, *p* < 0.0001; Syt1^DN^: n=19 NMJs from 5 larvae, *p* < 0.0001). **C-E**. Quantification of AZ sum Cac fluorescence (**C**, Ctrl: n=18 NMJs from 5 larvae; TeNT: n=19 NMJs from 5 larvae, *p* < 0.0001; Syt1^DN^: n=19 NMJs from 5 larvae, *p* < 0.0001), BRP (**D,** Ctrl: n=18 NMJs from 5 larvae; TeNT: n=19 NMJs from 5 larvae, *p* < 0.0001; Syt1^DN^: n=19 NMJs from 5 larvae, *p* < 0.0001) and RBP (**E**, Ctrl: n=18 NMJs from 5 larvae; TeNT: n=19 NMJs from 5 larvae, *p* < 0.0001; Syt1^DN^: n=19 NMJs from 5 larvae, *p* < 0.0001) across the 3 genotypes. **F.** Representative images of Cac, BRP and RBP immunolabelling within synaptic boutons of 3^rd^ instar M1 NMJs of controls (left panel) or following expression of *rab3* RNAi alone (2^nd^ panel) or co-expressed with TeNT (3^rd^ panel) or Syt1^DN^ (right panel) with the MN1-Ib-Gal4 driver. **G.** Quantification of AZ Cac area in the four genotypes (Ctrl: n=19 NMJs from 5 larvae*; rab3* RNAi: n=19 NMJs from 5 larvae, *p* = 0.0002; *rab3* RNAi + TeNT: n=19 NMJs from 5 larvae, *p* < 0.0001; *rab3* RNAi + Syt1^DN^: n=19 NMJs from 5 larvae, *p* < 0.0001). **H-J.** Quantification of AZ sum fluorescence for Cac (**H**, Ctrl: n=19 NMJs from 5 larvae*; rab3* RNAi: n=19 NMJs from 5 larvae, *p* = 0.019; *rab3* RNAi + TeNT: n=19 NMJs from 5 larvae, *p* < 0.0001; *rab3* RNAi + Syt1^DN^: n=19 NMJs from 5 larvae, *p* < 0.0001), BRP (**I,** Ctrl: n=19 NMJs from 5 larvae*; rab3* RNAi: n=19 NMJs from 5 larvae, *p* = 0.0009; *rab3* RNAi + TeNT: n=19 NMJs from 5 larvae, *p* < 0.0001; *rab3* RNAi + Syt1^DN^: n=19 NMJs from 5 larvae, *p* < 0.0001) and RBP (**J**, Ctrl: n=19 NMJs from 5 larvae*; rab3* RNAi: n=19 NMJs from 5 larvae, *p* = 0.0009; *rab3* RNAi + TeNT: n=19 NMJs from 5 larvae, *p* < 0.0001; *rab3* RNAi + Syt1^DN^: n=19 NMJs from 5 larvae, *p* < 0.0001) across the 4 genotypes. **K.** Representative images of Cac and HRP immunolabelling within synaptic boutons of M1 NMJs of controls or following expression of *rab3* RNAi, Syt1^DN^ or TeNT alone, or *rab3* RNAi co-expressed with TeNT or Syt1^DN^, with the MN1-Ib-Gal4 driver. **L.** Quantification of AZ number normalized to NMJ HRP area across the six genotypes (Ctrl: n=22 NMJs from 5 larvae*; rab3* RNAi: n=11 NMJs from 4 larvae, *p* < 0.0001; TeNT: n=21 NMJs from 5 larvae, *p* < 0.0001; Syt1^DN^: n=19 NMJs from 5 larvae, *p* < 0.0001; *rab3* RNAi + TeNT: n=15 NMJs from 4 larvae, *p* < 0.0001; *rab3* RNAi + Syt1^DN^: n=24 NMJs from 5 larvae, *p* < 0.0001). Statistical significance determined with one-way ANOVA followed by Tukey’s multiple comparisons test. Asterisks denote p-values of: *, p ≤ 0.05; **, p ≤ 0.01; ***, p ≤ 0.001). Absolute values and individual statistical tests are summarized in Table S1.

To determine if the increase in AZ size following activity reduction requires the Rab3 pathway, UAS-*rab3* RNAi constructs were co-expressed in MN1-Ib with either TeNT or Syt1^DN^. Expression of *rab3* RNAi alone induced synaptic defects at M1 NMJs similar to *rab3* mutants, including increased AZ area (Fig. 8F, G), enhanced BRP, RBP and Cac levels at these AZs (Fig. 8H-J), and decreased AZ density (Fig. 8L). Expression of TeNT or Syt1^DN^ together with *rab3* RNAi led to a synergistic enlargement of AZ size (Fig. 8F, G) and even higher levels of BRP, RBP and Cac accumulation compared to controls or expression of TeNT, Syt1^DN^ or *rab3* RNAi alone (Fig. 8H-J). Beyond the increase in AZ size and protein content, expression of TeNT or Syt1^DN^ at M1 NMJs also resulted in a reduction in AZ number (Fig. 8L), indicating neuronal silencing decreases AZ seeding. Co-expression of *rab3* RNAi with TeNT or Syt1^DN^ resulted in a synergistic decrease in AZ density as well (Fig. 8L).

To reduce synaptic activity independent of a blockage in SV fusion by TeNT and Syt1^DN^, UAS-RNAi knockdown of the Para voltage-gated Na^+^ channel was performed using the MN1-Ib Gal4 driver. Like TeNT and Syt1^DN^, blocking action potential generation in MN1-Ib increased AZ Cac area and Cac accumulation (Supplemental Fig. 2A-C). These effects were also enhanced by co-expression of *rab3* RNAi. Given loss of Para disrupts action-potential triggered evoked release and not spontaneous fusion, the effects of activity reduction on AZ maturation and size are likely to be downstream of signaling pathways primarily associated with evoked output. Consistent with evoked release being the major driver for AZ size expansion, the Syt1^DN^ transgene dramatically reduces evoked output, but actually increases spontaneous release rate^35^. Together, these data indicate synaptic inactivity and loss of Rab3 trigger AZ enlargement via distinct mechanisms, with neither pathway alone reaching an upper limit for these processes.

To examine how synaptic inactivity decreases AZ number, we used CRISPR to endogenously tag Unc13A and Unc13B splice variants in the same transgenic line to directly visualize early and late AZ scaffold distribution *in vivo* (Fig. 9). The Unc13A exon was tagged with mRuby and the Unc13B exon was tagged with mClover. The density of AZs containing the late scaffold protein Unc13A was reduced at 3^rd^ instar M1 NMJs in *rab3* mutants (2.4-fold) and larvae expressing either TeNT (1.9-fold) or Syt1^DN^ (1.8-fold) with the MN1-Ib driver (Fig. 9A, C). In contrast, the density of AZs containing the early scaffold Unc13B was similar between controls and *rab3* mutants, while expression of TeNT (1.8-fold) or Syt1^DN^ (1.8-fold) reduced Unc13B^+^ AZs to a similar extent to Unc13A^+^ sites (Fig. 9A-D). We conclude that unlike Rab3 that controls the distribution of late scaffolds across the AZ population, synaptic inactivity results in a global reduction in AZ seeding that affects both early and late AZ components.

**Fig. 9.**
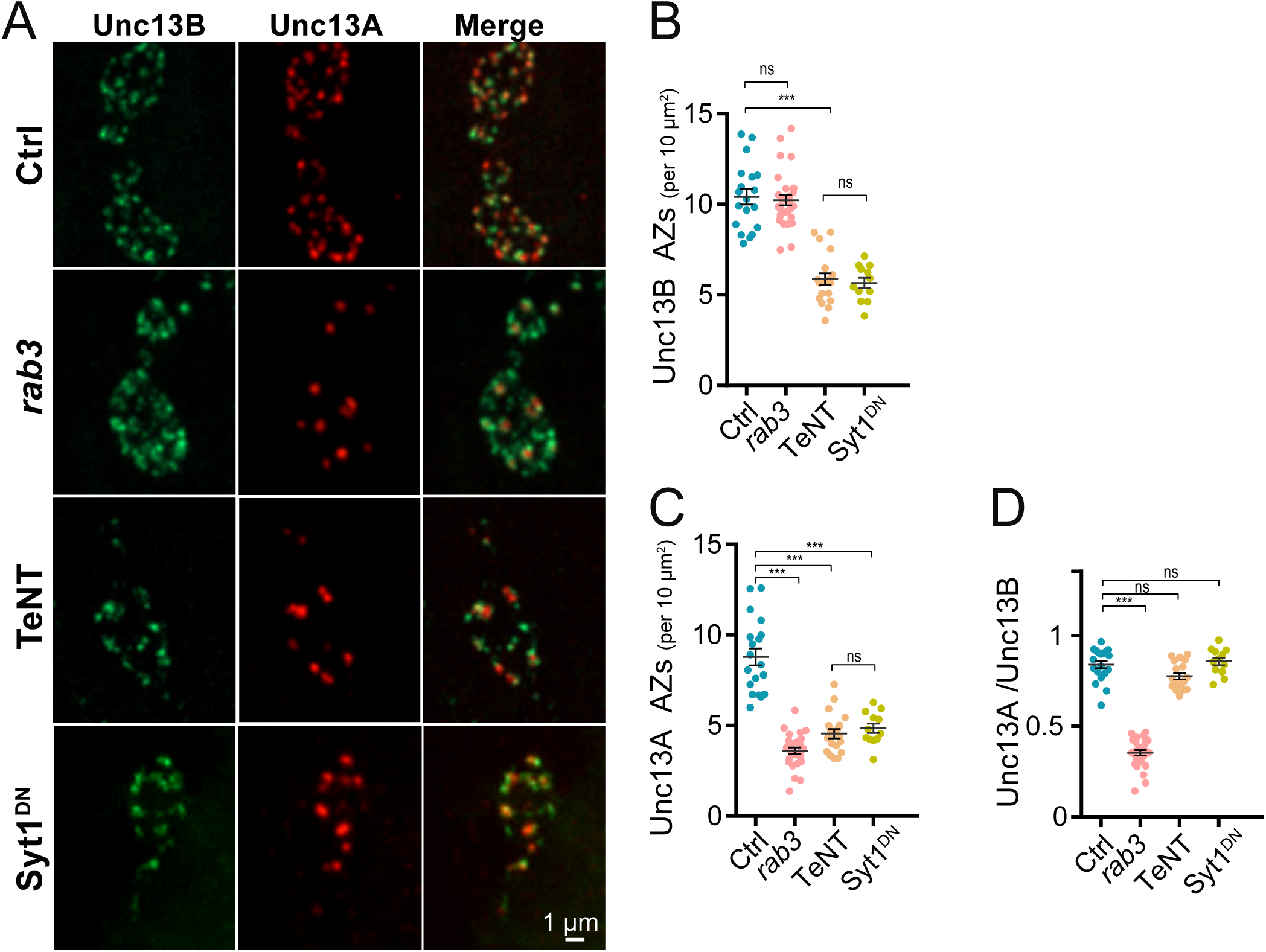
Presynaptic silencing reduces seeding of both early and late AZ scaffolds. **A.** Representative images of Unc13B-mClover and Unc13A-mRuby localization within synaptic boutons of 3^rd^ instar M1 NMJs of controls (top panel) or following expression of *rab3* RNAi (2^nd^ panel), TeNT (3^rd^ panel) or Syt1^DN^ (bottom panel) with the MN1-Ib-Gal4 driver. The merged image is shown on the right. Scale bar = 1 µm. **B**. Quantification of Unc13B^+^ AZ number (per 10 μm^2^) across the four genotypes (Ctrl: n=19 NMJs from 5 larvae*; rab3* RNAi: n=28 NMJs from 6 larvae, *p* = 0.99; TeNT: n=19 NMJs from 5 larvae, *p* < 0.0001; Syt1^DN^: n=12 NMJs from 4 larvae, *p* < 0.0001). **C**. Quantification of Unc13A^+^ AZ number (per 10 μm^2^) across the four genotypes (Ctrl: n=19 NMJs from 5 larvae*; rab3* RNAi: n=28 NMJs from 6 larvae, *p* < 0.0001; TeNT: n=19 NMJs from 5 larvae, *p* < 0.0001; Syt1^DN^: n=12 NMJs from 4 larvae, *p* < 0.0001). **D**. Quantification of the Unc13A^+^/Unc13B^+^ AZ ratio across the four genotypes (Ctrl: n=19 NMJs from 5 larvae*; rab3* RNAi: n=28 NMJs from 6 larvae, *p* < 0.0001; TeNT: n=19 NMJs from 5 larvae, *p* = 0.07; Syt1^DN^: n=12 NMJs from 4 larvae, *p* = 0.57). Statistical significance determined with one-way ANOVA followed by Tukey’s multiple comparisons test. Asterisks denote p-values of: ***, p ≤ 0.001. Absolute values and individual statistical tests are summarized in Table S1.

### Synaptic inactivity increases T-bar area while loss of Rab3 causes multiple T-bars per AZ

Transmission electron microscopy (TEM) of *Drosophila* synapses reveals presynaptic AZs are highlighted by a central electron dense body referred to as the T-bar^37^. Most AZs contain a single T-bar composed of scaffolding proteins like BRP^11,38^, suggesting a modular organization for these release sites. To compare how loss of Rab3 and synaptic inactivation alter the ultrastructure of individual release sites, TEM was performed at larval M1 NMJs from controls or lines expressing TeNT, *rab3* RNAi or both using the MN1-Ib driver (Fig. 10A). Silencing neuronal activity resulted in a 39% increase in the size of individual T-bars (Fig. 10B, measured as length of the upper T-bar platform), but did not cause increased incorporation of T-bars at single AZs in MN1-Ib NMJs (Fig. 10C). Loss of Rab3 resulted in a distinct phenotype, with no change in individual T-bar size but a ∼1.5-fold increase in the number of T-bars per AZ, as previously observed^22^. Reducing activity together with Rab3 depletion resulted in more severe AZ disorganization, with multiple enlarged T-bars per release site (Fig. 10A-C). In contrast to the increase in T-bar size or number, the length of the electron dense synaptic cleft was only increased following TeNT expression in *rab3* knockdown larvae (Fig. 10D).

**Fig. 10.**
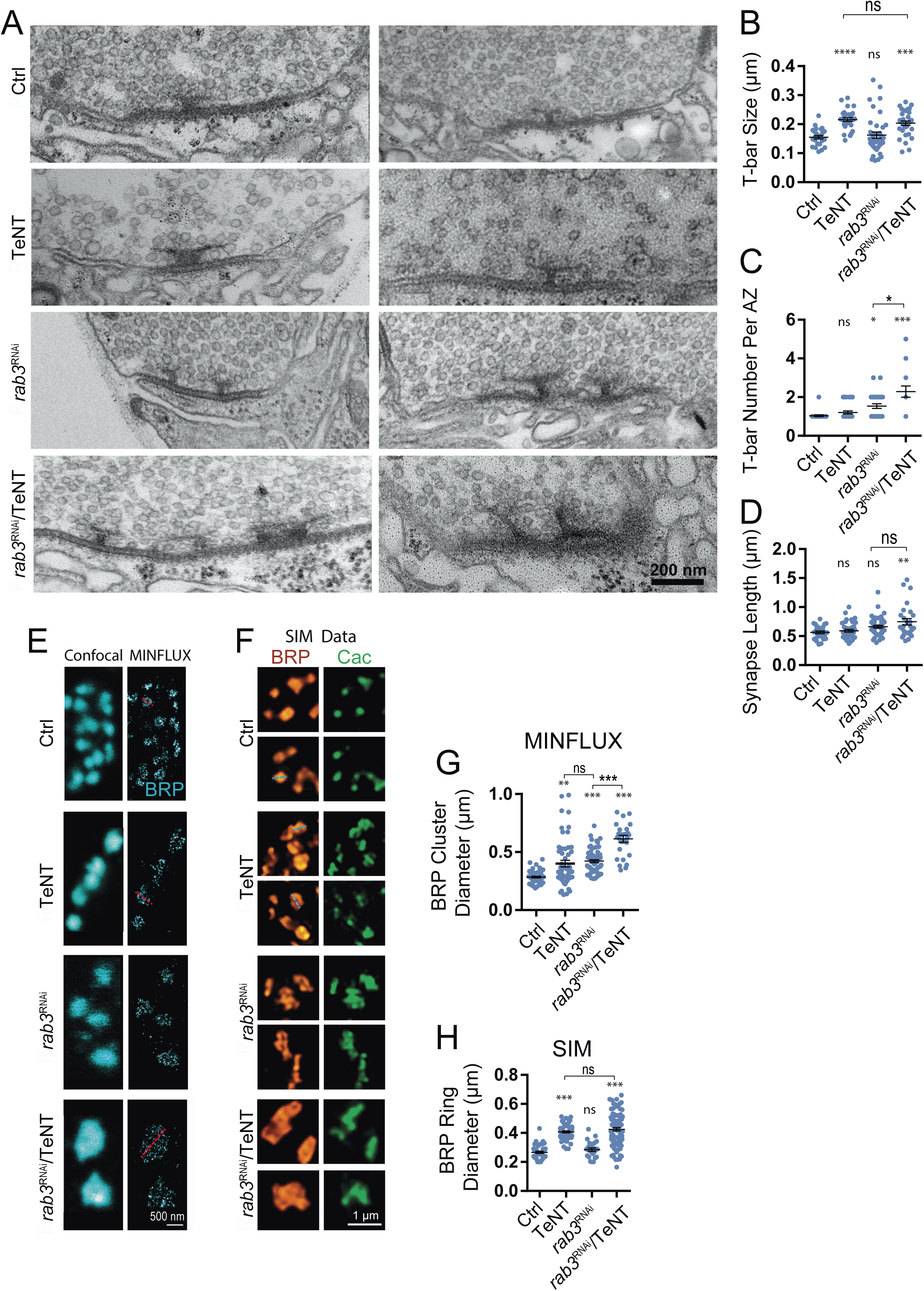
Presynaptic silencing increases T-bar area while *rab3* mutants contain multiple T-bars per AZ. **A.** Representative EM images of synaptic boutons at 3^rd^ instar M1 NMJs of controls (top panel) or following expression of TeNT (2^nd^ panel), *rab3* RNAi (3^rd^ panel) or *rab3* RNAi + TeNT (bottom panel) with the MN1-Ib-Gal4 driver. Scale bar = 200 nm. **B**. Quantification of T-bar size (length of the upper T-bar platform) across the four genotypes (Ctrl: n=25 T-bars from 3 larvae; TeNT: n=30 T-bars from 3 larvae, *p* < 0.0001; *rab3* RNAi: n=39 T-bars from 3 larvae, *p* = 0.88; *rab3* RNAi + TeNT: n=33 T-bars from 3 larvae, *p* = 0.0007). **C**. Quantification of T-bar number per AZ across the four genotypes (Ctrl: n=25 AZs from 3 larvae; TeNT: n=29 AZs from 3 larvae, *p* = 0.74; *rab3* RNAi: n=28 AZs from 3 larvae, *p* = 0.04; *rab3* RNAi + TeNT: n=21 AZs from 3 larvae, *p* < 0.0001). **D**. Quantification of synapse length (measurement of synaptic cleft electron dense material length) across the four genotypes (Ctrl: n=26 synapses from 3 larvae; TeNT: n=37 synapses from 3 larvae, *p* = 0.90; *rab3* RNAi: n=45 synapses from 3 larvae, *p* = 0.09; *rab3* RNAi + TeNT: n=27 synapses from 3 larvae, *p* = 0.001). **E**. Representative confocal and MINFLUX images of synaptic boutons at 3^rd^ instar M1 NMJs of controls (top panel) or following expression of TeNT (2^nd^ panel), *rab3* RNAi (3^rd^ panel) or *rab3* RNAi + TeNT (bottom panel) with the MN1-Ib-Gal4 driver. The red dashed line denotes a representative BRP cluster diameter measurement. Scale bar = 500 nm. **F**. Representative SIM images of synaptic boutons at 3^rd^ instar M1 NMJs of controls (top panel) or following expression of TeNT (2^nd^ panel), *rab3* RNAi (3^rd^ panel) or *rab3* RNAi + TeNT (bottom panel) with the MN1-Ib-Gal4 driver. The blue dashed line denotes a representative BRP ring diameter measurement. Scale bar = 1 μm. **G**. Quantification of BRP cluster diameter measured with MINFLUX across the four genotypes (Ctrl: n=49 AZs from 3 larvae; TeNT: n=53 AZs from 3 larvae, *p* = 0.0001; *rab3* RNAi: n=65 AZs from 3 larvae, *p* < 0.0001; *rab3* RNAi + TeNT: n=28 AZs from 3 larvae, *p* < 0.0001). **H**. Quantification of BRP ring diameter measured with SIM across the four genotypes (Ctrl: n=45 AZs from 3 larvae; TeNT: n=59 AZs from 3 larvae, *p* < 0.0001; *rab3* RNAi: n=24 AZs from 3 larvae, *p* = 0.71; *rab3* RNAi + TeNT: n=95 AZs from 3 larvae, *p* < 0.0001). Statistical significance determined with one-way ANOVA followed by Tukey’s multiple comparisons test. Asterisks denote p-values of: *, p ≤ 0.05; **, p ≤ 0.01; ***, p ≤ 0.001. Absolute values and individual statistical tests are summarized in Table S1.

To examine T-bar morphology following activity reduction at the nanoscale, MINFLUX imaging^39^ with a resolution comparable to EM was used, allowing visualization of filamentous BRP clusters at AZs (Fig. 10E). This single molecule localization technology determines initially unknown fluorophore positions by relating it to the know zero-intensity position of a donut-shaped excitation beam. The diameter of BRP clusters increased following TeNT or *Rab3* RNAi expression and was further enlarged when TeNT and *Rab3* RNAi were co-expressed (Fig. 10 E, G). Structural illumination microscopy (SIM) provided an optimal approach to image individual BRP AZ rings in larvae (Fig. 10F). Expression of TeNT resulted in a 53% enlargement in BRP ring diameter, while Rab3 reduction did not alter ring diameter but displayed multiple BRP rings at single AZs (Fig. 10H) as observed by TEM. When TeNT and *Rab3* RNAi were combined, larger BRP rings were present as clusters within individual AZs (Fig. 10F, H). These data indicate activity reduction alters AZ size via a morphologically distinct process compared to loss of Rab3.

To examine if increased AZ size caused by reduced presynaptic output may utilize a similar pathway as presynaptic homoeostatic plasticity (PHP) downstream of loss or blockage of GluRIIA^4,19,33^, Syt1^DN^ was expressed in MN1-Ib in null mutants lacking GluRIIA. In controls, AZ Cac area was indistinguishable at M1 and M9 NMJs, resulting in a M1/M9 AZ area ratio of ∼1 (Fig. 11A, B). While expression of Syt1^DN^ in MN1-Ib triggered AZ enlargement and a larger M1/M9 AZ area in heterozygous *GluRIIA^+/-^* controls, expression in *GluRIIA* homozygous mutants failed to induce an enlargement in AZ Cac area (Fig. 11B, C). In contrast, loss of GluRIIA did not prevent AZ size expansion in *rab3* mutants (Fig. 11A-C), consistent with distinct mechanisms mediating AZ enlargement as described above. Although GluRIIA is necessary for AZ enlargement following reduced presynaptic output, it was not required for the corresponding reduction in AZ number induced by Syt1^DN^ expression (Fig.11D). These data suggest increased AZ size is not strictly due to enhanced delivery of material over a reduced number of AZs. To determine if loss of GluRIIA alone resulted in an increase in AZ area that would preclude further increases following Syt1^DN^ expression, AZ Cac area was compared between controls and *GluRIIA* null mutants. No increase in AZ Cac area was observed in *GluRIIA* mutants alone (Fig. 11C), suggesting mechanisms mediating PHP-induced AZ remodeling diverge from those required for presynaptic silencing.

**Fig. 11.**
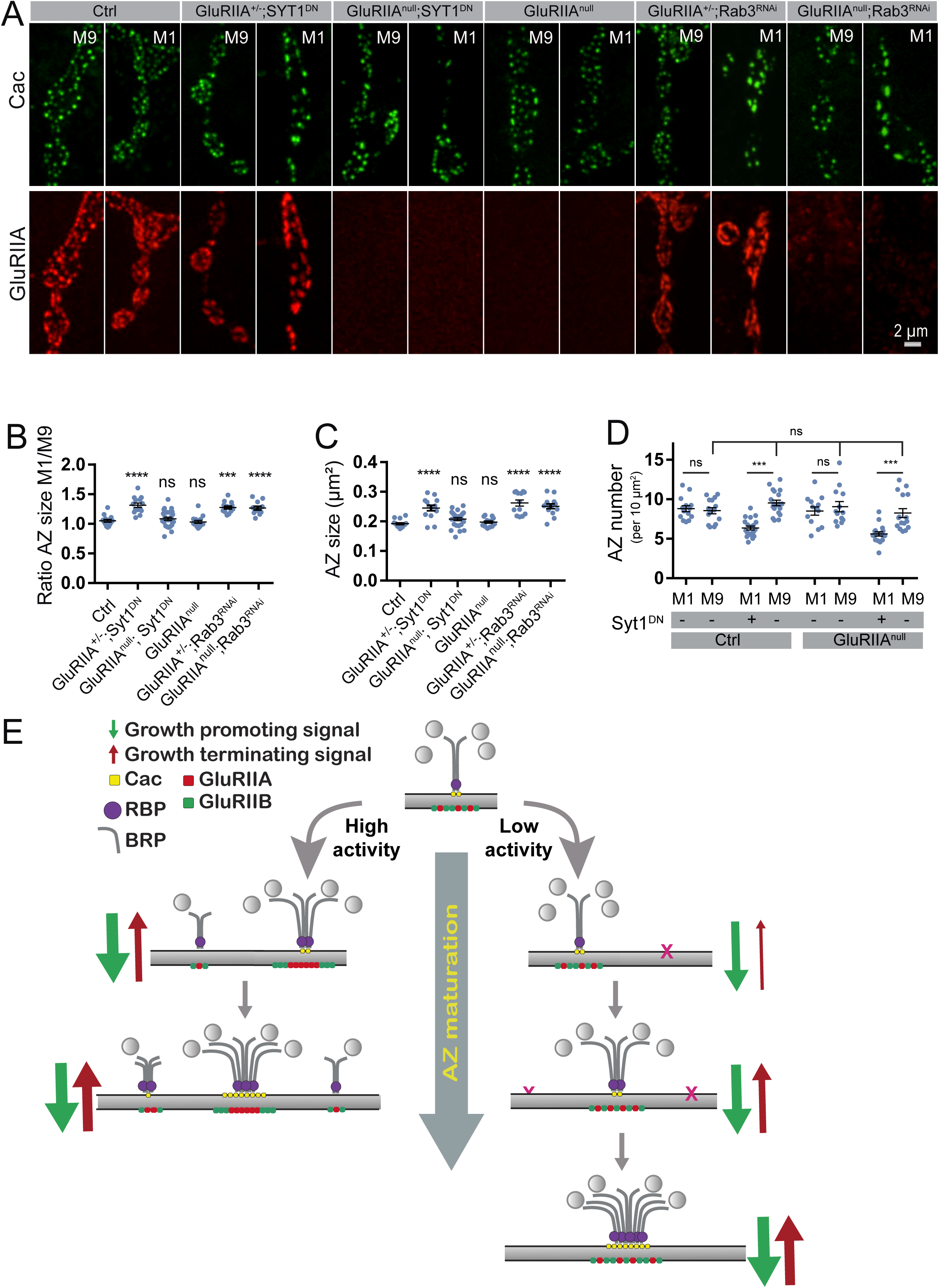
Loss of GluRIIA suppresses AZ enlargement following presynaptic silencing. **A.** Representative images of Cac (top panel) and GluRIIA (lower panel) immunolabelling at synaptic boutons of 3^rd^ instar M9 and M1 NMJs of controls, *GluRIIA^-/-^* null mutants or following expression of Syt1^DN^ or *rab3* RNAi with the MN1-Ib-Gal4 driver in control GluRIIA^+/-^ heterozygotes or GluRIIA^-/-^ nulls. **B**. Quantification of the M1/M9 ratio for AZ Cac area across the six genotypes (Ctrl: n=14 NMJs from 4 larvae; GluRIIA^+/-^ + Syt1^DN^: n=14 NMJs from 4 larvae, *p* < 0.0001; GluRIIA^-/-^ + Syt1^DN^: n=26 NMJs from 6 larvae, *p* = 0.84; GluRIIA^-/-^: n=14 NMJs from 4 larvae, *p* = 0.83; GluRIIA^+/-^ + *rab3* RNAi: n=13 NMJs from 4 larvae, *p* = 0.0002; GluRIIA^-/-^ + *rab3* RNAi: n=14 NMJs from 4 larvae, *p* < 0.0001). **C**. Quantification of AZ Cac area at M1 NMJs across the six genotypes (Ctrl: n=14 NMJs from 4 larvae; GluRIIA^+/-^ + Syt1^DN^: n=14 NMJs from 6 larvae, *p* < 0.0001; GluRIIA^-/-^ + Syt1^DN^: n=26 NMJs from 4 larvae, *p* = 0.30; GluRIIA^-/-^: n=14 NMJs from 4 larvae, *p* = 0.98; GluRIIA^+/-^ + *rab3* RNAi: n=13 NMJs from 4 larvae, *p* < 0.0001; GluRIIA^-/-^ + *rab3* RNAi: n=14 NMJs from 4 larvae, *p* < 0.0001). **D**. Quantification of M1 and M9 AZ number in controls or *GluRIIA^-/-^* null mutants with or without expression of Syt1^DN^ using the MN1-Ib-Gal4 driver (Ctrl M1: n=14 NMJs from 4 larvae; Ctrl M9: n=15 NMJs from 4 larvae, *p* = 0.99; M1 in MN1-Ib-Gal4>Syt1^DN^: n=20 NMJs from 5 larvae, *p* = 0.0003; M9 in MN1-Ib-Gal4>Syt1^DN^: n=19 NMJs from 5 larvae, *p* = 0.7; M1 in GluRIIA^-/-^: n=13 NMJs from 4 larvae, *p* = 0.99; M9 in GluRIIA^-/-^: n=13 NMJs from 4 larvae, *p* = 0.99; M1 in GluRIIA^-/-^ + MN1-Ib-Gal4>Syt1^DN^: n=17 NMJs from 4 larvae, *p* < 0.0001; M9 in GluRIIA^-/-^ + MN1-Ib-Gal4>Syt1^DN^: n=16 NMJs from 4 larvae, *p* = 0.90). Statistical significance determined with one-way ANOVA followed by Tukey’s multiple comparisons test. Asterisks denote p-values of: *, p ≤ 0.05; **, p ≤ 0.01; ***, p ≤ 0.001. Absolute values and individual statistical tests are summarized in Table S1. **E**. Model for the role of presynaptic output in synapse maturation. Under high activity conditions, both AZ and PSD maturation rate is enhanced, and more synapses are seeded. Once AZs reach their normal high *P_r_* state, a maturation terminating signal requiring postsynaptic GluRIIA inhibits further AZ enlargement. Under conditions where presynaptic output is dramatically reduced, AZ and PSD maturation is delayed, and fewer synapses form. The reduction in presynaptic output also disrupts the normal maturation stop signal, resulting in larger AZs containing more Cac and scaffolding proteins.

## Discussion

In the current study we describe how AZ maturation in *Drosophila* larval MNs controls their output strength, and how this process is regulated by neuronal activity. Prior quantal imaging studies at *Drosophila* NMJs demonstrated a ∼50-fold heterogeneity in *P_r_* across the hundreds of AZs formed by a single neuron^7,12,13,18,40–42^. One model for this heterogeneity predicts each AZ is fated to have a distinct *P_r_* that is maintained over the life of the AZ. This could arise from AZ components being delivered in variable abundance and/or composition that are deposited together at forming synapses, or through the presence of unique components that direct future output to a defined set point. An alternative model is that AZs are in the process of maturing to a high *P_r_* state, with heterogeneity reflecting distinct stages of maturation across the AZ population. Using mMaple-tagged GluRs to timestamp individual synapses, we find that older AZs have a higher evoked *P_r_* than newly formed release sites. In addition, we observe >85% of newly formed synapses will accumulate BRP within several days, indicating the majority of AZs are progressing to a higher *P_r_* state. Together with the sequential addition of specific AZ proteins observed *in vivo*, these data indicate AZs formed earlier in development have more time to accumulate scaffolding proteins and Ca^2+^ channels that promote higher *P_r_* compared to newly formed AZs. By timestamping synapses in the 1^st^ instar stage and recording their output in late 3^rd^ instars, early formed AZs also stably maintain their high *P_r_* state across the 6-day developmental window of larval development.

Supporting a developmental AZ maturation model at larval NMJs, previous studies using serial intravital imaging identified sequential addition of early versus late AZ scaffolding proteins^3,6,11,43–45^. Using fixed and live imaging of a large panel of AZ proteins in the current study, Liprin-α is the first arriving AZ component detected, followed by the early scaffold proteins Syd1, RIM, and Unc13B. BRP, RBP and Unc13A scaffolds appear later and around the same time, with Cac channels arriving several hours after RBP. In addition, the abundance of early AZ scaffolding proteins correlates poorly with Cac levels and evoked *P_r_* compared to late AZ scaffolds. We also find that AZ maturation state regulates action-potential independent spontaneous SV release. Although AZs lacking Cac channels would not be expected to participate in evoked release, we observe many of these sites support spontaneous fusion. We propose “spontaneous-only” AZs at *Drosophila* NMJs^13^ largely reflect an earlier stage of maturation prior to accumulation of sufficient Ca^2+^ channels to drive evoked fusion, rather than a unique AZ state dedicated to this mode of synaptic communication. As such, AZ maturation regulates both release mode and output strength for individual synaptic sites. It will be interesting to determine if long-lived adult AZs that require more extensive protein turnover stably maintain a high *P_r_* state after maturation or instead have a “lifetime” and are eventually replaced.

How the sequential addition of AZs proteins is regulated *in vivo* within synaptic boutons is unclear. Prior evidence suggests early and late scaffolds can be assembled on a precursor vesicle from the trans-Golgi and transported together along axons^25,46–48^. Our data suggest that if AZ components are co-delivered in a common transport vesicle, they would need to be disassembled within synaptic boutons and sequentially added as AZs mature. Additionally, new AZs are continually forming at NMJs throughout development, with a ∼1.7-fold increase in AZ number each day during early larval stages. This rapid addition of new release sites suggests the time course for AZ maturation could vary across development. Only a small number of AZs (∼8-10) are initially formed at embryonic NMJs^7^, and we hypothesize enough material and assembly machinery are available for these sites to mature more quickly. With each day of development, a near doubling of AZ number occurs (for example: 10 AZs day one, 20 AZs day two, 40 AZs day three, 80 AZs day four, etc). When many more AZs are being added, shortages of material and/or regulatory components to rapidly assemble this large AZ cohort could delay maturation rate. We suggest this mechanism accounts for the large population of low *P_r_* AZs observed at the late 3^rd^ instar stage when many newly added sites have not had sufficient time to fully mature. Given larval MNs burst fire at 20-30 Hz^49^, having high *P_r_* AZs undergoing depression and low *P_r_* AZs undergoing facilitation is likely beneficial for providing a mechanism to maintain release output across the AZ population during high-frequency firing^40^.

Neuronal activity provides an additional mechanism to regulate synapse maturation. Indeed, prior studies found elevated neuronal activity at *Drosophila* NMJs can enhance synaptic growth^31,50–52^ and promote faster maturation of postsynaptic GluR fields^7,53–56^. In the current study, we find changes in neuronal activity in either direction influence synapse formation and maturation. Disrupting SV fusion with TeNT or Syt1^DN^, blocking action potentials by knocking down the voltage-gated Na^+^ channel Para, or reducing release in CRISPR-deleted Cac NMJs results in slower material accumulation but ultimately an enlargement in both pre and postsynaptic compartments. The expansion in AZ size and protein content is accompanied by a reduction in AZ number, suggesting the absence of presynaptic output decreases seeding of new AZs and promotes insertion of AZ material into preexisting release sites.

The mechanisms by which silenced presynaptic neurons control AZ size versus AZ addition are likely to be executed by distinct molecular programs, as *GluRIIA* mutants fully suppress AZ expansion but do not prevent the reduction in AZ number. We hypothesize AZ expansion requires the postsynaptic muscle to report reduced neurotransmission through a GluRIIA-mediated retrograde signal, while AZ seeding may instead represent a cell autonomous mechanism or function via a GluRIIA-independent postsynaptic signal. We observe a strong synergistic effect on increased AZ size and protein accumulation between the Rab3-dependent pathway and neuronal silencing, suggesting these represent independent mechanisms for controlling synapse size. Indeed, EM analysis revealed increased T-bar diameter in silenced neurons versus multiple normal size T-bars per AZ in *rab3* mutants. Using dual CRISPR-tagging of the two splicing isoforms of Unc13 (the early Unc13B scaffold and the late Unc13A scaffold), we find *rab3* mutants and neuronal silencing also decrease the number of functional release sites through independent mechanisms. While *rab3* mutants terminate AZ formation at later stages (reduced Unc13A^+^ AZs), the ability to seed AZs containing early scaffolds remains intact (normal Unc13B^+^ AZ number). In contrast, neuronal silencing reduces the number of both Unc13A^+^ and Unc13B^+^ AZs, indicating a mechanism that targets the earliest steps of AZ seeding.

On the postsynaptic side, the levels of GluRIIA are reduced at PSDs opposed to silenced AZs early in development. In addition, the normal segregation of GluRIIA into the PSD center is slowed, leading to a larger PSD area containing the receptors. Altered GluRIIA incorporation into PSDs following TeNT expression has been previously observed at *Drosophila* NMJs^15^, suggesting several aspects of GluRIIA dynamics are sensitive to synaptic activity. Using PC of mMaple-tagged GluRIIE, we find GluRs are surprisingly stable once inserted into a PSD, with little turnover and no lateral movement between neighboring PSDs. Consistent with these observations, studies using FRAP of fluorescently-tagged GluRIIA observed reduced recovery at mature PSDs, while newer GluRs readily accumulated at younger PSDs^15^. Together, these data indicate modifications (i.e. phosphorylation) that alter GluR function at *Drosophila* NMJs will have effects limited to PSDs that contains the affected receptors, consistent with the distinct effects on GluR accumulation at BRP^+^ versus BRP^-^ AZs in *rab3* mutants. Given GluRs are essentially immobilized at mature PSDs, this stability also contributes to the enhanced function of older synapses that contain more GluRs by promoting a larger postsynaptic response to released neurotransmitters.

The effects of activity on synapse maturation are not limited to *Drosophila*, as a lack of activity in cultured mammalian spinal neurons that were silenced display increased AMPA receptor accumulation^57^. Likewise, silencing of activity in hippocampal neurons alters the nanoscale structure of the AZ matrix downstream of changes in the local actin cytoskeleton, resulting in enrichment of AZ components including Ca^2+^ channels^58^. Enhancement in the levels of presynaptic Ca^2+^ channels in response to activity silencing has also been observed in mammalian cortical neuron cultures^59^. Although the molecular pathways by which activity regulates AZ seeding and maturation will require further work, our findings suggest a model to drive further experimental analysis (Fig. 11E). In the early steps of AZ development, enhanced synaptic activity has a positive role in promoting AZ material accumulation and seeding of new AZs. Once an individual AZ reaches a preferred state of output, GluRIIA receptors in the opposed PSD facilitate transmission of a local retrograde signal that terminates further AZ expansion. In cases where presynaptic AZs are silenced, material accumulation is slowed but the signal to inhibit AZ expansion is reduced, resulting in larger AZs that contain more presynaptic material by the end of development. A separate pathway downstream of reduced neuronal activity would reduce seeding of new AZs as well. A diverse set of pathways, including BMPs, Wingless, FGFs, and neurotrophins, have been previously implicated in NMJ development and AZ morphology in *Drosophila*^2,60^. In addition, multiple trans-synaptic cell adhesion molecules (CAMs), including neurexin–neuroligin, cadherins, and teneurins, function in synaptic development and transsynaptic signaling^61^. Finally, proteins required for enhanced presynaptic output during PHP downstream of GluRIIA inhibition^62^ could also that alter AZ maturation downstream of presynaptic silencing. In summary, our data indicate AZs require a developmental maturation process at *Drosophila* NMJs that controls both release mode and output strength. Alterations in neuronal activity alter AZ maturation, with neuronal silencing resulting in expansion of AZ size with excess material accumulation at the expense of AZ number.

## Materials and methods

### Drosophila stocks

*Drosophila melanogaster* were cultured on standard medium at 25°C. Male larvae were used for all experiments to facilitate crossing as fluorescently labeled Cac is located on the X chromosome. The syt1^DN^ line used in this study was UAS-Syt1^C2BD356N,D362N 35^. GluRIIB-GFP, GluRIIA-RFP, UAS-Liprin-α-mStrawberry, UAS-Syd1-mStrawberry, UAS-RBP-mCherry and UAS-Rim-mCherry were provided by Stephan Sigrist. The *rab3* null mutant (*rab3^rup^*) was provided by Ethan Graf. Other lines used in the study include UAS-TeNT (BDSC #28837), sgRNA for Cac (BDSC #85863), sgRNA for Para (BDSC #33923), RNAi for Rab3 (VDRC #330151), Ib-specific Gal4 GMR94G06 (BDSC #40701), endogenously CRISPR-tagged cac-GFP^19^, and UAS-cas9^63^. For AZ *P_r_* mapping, 44H10-LexA (provided by Gerry Rubin), LexAOp-myr-jGCaMP7s^17^, and endogenously CRISPR-tagged Cac^TagRFP^ ^19^ transgenic lines were used. The GluRIIA null mutant used contains a CRISRP-induced frame shift (provided by Dion Dickman)^64^.

### Generation of GluRIIE-mMaple and fluorescently labeled Unc13A/B lines

To generate GluRIIE mMaple, GluRIIE sequence was obtained from plasmid FI04462 (DGRC stock 1621963) and mMaple was amplified from a plasmid (Addgene #141151). The two constructs were cloned into pBid-UASc (Addgene #35200) using EcoRI and XbaI. To create CRISPR-tagged Unc13A, four guide RNAs (gRNAs) were selected using the CRISPR Optimal Target Finder^65^: gRNA1 AGCTCGGCAACGATGGCATT gRNA2 CATGCAGGTGTTACGCCAAA gRNA3 AAAAAAAAAACGCTCTTGAG gRNA4 CCTGCCCTCAATTAAAGTAG

These gRNAs were fused with the pCFD5 expression vector (Addgene #73914) according to the Gibson assembly protocol using NEBuilder HighFidelity DNA Assembly Cloning Kit (E5520). mRuby was amplified from a plasmid (Addgene #105802) and flanked with right and left homology arms corresponding to the end of the first exon for Unc13A. Small flex regions were added before (ggcggaagc) and after (ggaggcagt) the mRuby sequence. A similar strategy was used to generate Unc13B tagged with mClover. Four gRNAs were assembled in pCFD5, targeting the area close to the end of the first exon corresponding to Unc13B: gRNA1 GTAGGCAATGAAACCGCTGT gRNA2 AACCCTTCCGAACCGATTTA gRNA3 AACCAGGGCTGTGAACCATT gRNA4 AAGATGAAATGATAAAGGAC

The mClover sequence was copied from a plasmid (Addgene #105778) and flanked to right and left homology arms corresponding to an 800 base pair area close to the end of the first exon. These constructs were co-injected by BestGene Inc. (Chino Hills, CA, USA) into vas-Cas9 embryos (BDSC #51324) and positive transformants were selected by PCR amplification of sequences corresponding to mRuby and mClover.

### mMaple photoconversion

Larvae were collected, briefly washed in HL3 saline, and placed ventral side up on a glass slide with a drop of halocarbon oil. To restrict larval movement, a coverglass was placed on top of larvae and attached with plastalina clay. Photoconversion was performed using a 10X Zeiss Plan ApoChromat objective. Samples were illuminated using a Lumen Dynamics X-Cite XYLIS LED light source with a 360/51 BP filter for 15 seconds.

### Optical AZ *P_r_* mapping

3^rd^ instar larvae were dissected in Ca^2+^-free HL3 containing 20 mM MgCl. After dissection, preparations were maintained in HL3 with 20 mM MgCl and 1.0 mM Ca for 5 min. Optical AZ *P_r_* mapping was performed on a PerkinElmer system equipped with a spinning-disk confocal head (CSU-X1; Yokagawa, Japan) and Hamamatsu C9100–13 ImagEM EM CCD camera (Hamamatsu, Hamamatsu City, Japan) as previously described^7^. An Olympus LUMFL N 60 X objective with a 1.10 NA was used to acquire GCaMP7s imaging data at 8 Hz. A dual channel multiplane Z-stack at M4 NMJs in segments A3 - A4 was acquired at the beginning of each experiment to identify AZ position using expression of Cac^TagRFP^ or fluorescently-labeled AZ scaffolds (Liprin-α, Syd1, RBP or Unc13A). Single focal plane videos were then recorded while MNs were stimulated with a suction electrode at 1 Hz for 3-5 min. Slight z-drift was manually corrected during each imaging session, while imaging sessions with significant muscle movement were discarded. For experiments using time-stamped PSDs, UAS-GluRIIE-mMaple was expressed postsynaptically using Mef2-Gal4 (BDSC #27390). AZ or PSD position was re-imaged every 25 sec during constant video recording of myr-GCaMP7s. The dual channel stack was merged into a single plane using the max intensity projection algorithm from Volocity 6.5.0 software (Quorum Technologies Inc.). Images of all AZs or PSDs was added to the myr-GCaMP7s stimulation video and detected automatically using the spot finding function of Volocity. Equal size ROIs were assigned to each. In cases where the software failed to label visible AZs or PSDs, ROIs were added manually. GCaMP7s peak flashes were detected and assigned to ROIs based on centroid proximity with a 0.8 µm threshold. Evoked events were identified as frames with three or more simultaneous GCaMP events recorded at the NMJ. The time and location of events were imported into Excel for further analysis. Evoked GCaMP events per ROI were divided by the number of stimulations to calculate AZ *P_r_*.

### Confocal imaging and data analysis

Confocal images were acquired using a PerkinElmer system equipped with a spinning-disk confocal head (CSU-X1; Yokagawa, Japan) and Hamamatsu C9100–13 ImagEM EM CCD camera (Hamamatsu, Hamamatsu City, Japan). A Zeiss plan-APOCHROMAT 63× objective with 1.40 NA was used for imaging of stained preparations. A 3D image stack was acquired for each NMJ imaged. Image analysis was performed in Volocity 3D Image Analysis software (Quorum Technologies Inc.) and 3D stack images were merged into a single plane for 2D analysis using the max intensity projection algorithm from Volocity 6.5.0 software. Analysis of Cac, BRP, RBP, GluRIIB and GluRIIA intensities were performed via the ‘find object’ algorithm that detects fluorescent areas above a preset threshold. All images in the same dataset where tiled together and one algorithm of analysis was applied to each image. Immunoreactive proteins were imaged at segments A3 and A4 of M4 for all experiments unless indicated. For M1 specific manipulations, neighboring M9 NMJs in the same larvae were used as control.

### Immunocytochemistry

Larvae were dissected in HL3 solution and fixed in 4% paraformaldehyde for 10 min, washed in Ca^2+-^free HL3, and blocked and permeabilized for 1 hr in PBS containing 0.1% Triton X-100, 2.5% NGS, 2.5% BSA and 0.1% sodium azide. Samples were incubated overnight at 4°C in blocking solution containing primary antibodies and then washed for 1 hr in blocking solution. Samples were incubated for 1 hr at room temperature in blocking solution containing fluorophore-conjugated secondary antibodies. Primary antibodies used in this study were mouse anti-BRP at 1:500 (Nc82 DSHB, Iowa City, IA) and rabbit anti-RBP at 1:500 (provided by Stephan Sigrist). Secondary antibodies used in this study were goat anti-mouse Alexa Fluor 607-, 546-, or 488-conjugated anti-mouse IgG at 1:1000 (Invitrogen, #s A21237, A11030, and A32723) and goat anti-rabbit Alexa Fluor 647-conjugated IgG at 1:1000 (Molecular probes #4414). For HRP staining, samples were incubated in DyLight 649-conjugated HRP at 1:500 (#123-605-021; Jackson Immuno Research, West Grove, PA). Samples were mounted in Vectashield (Vector Laboratories, Burlingame, CA) before imaging.

### Electron microscopy

3^rd^ instar larvae were dissected in Ca^2+-^free HL3.1 solution and fixed in 1% glutaraldehyde, 4% formaldehyde, and 0.1 M sodium cacodylate for 10 min at room temperature. To identify specific muscle NMJs, cacti pins were positioned next to M1 and larvae were transferred into the glass vials. Fresh fixative was added and samples were incubated for 1 hr at room temperature. After washing samples in 0.1 M sodium cacodylate and 0.1 M sucrose, samples were stained for 30 min in 1% osmium tetroxide and 1.5% potassium ferrocyanide in 0.1 M sodium cacodylate solution. After washing with 0.1 M sodium cacodylate, samples were stained for 30 mins in 2% uranyl acetate and dehydrated through a graded series of ethanol and acetone, before embedding in epoxy resin (Embed 812; Electron Microscopy Sciences). Thin sections (50–60 nm) were collected on Formvar/carbon-coated copper slot grids and contrasted with lead citrate and uranyl acetate. Sections were imaged at 49,000 × magnification at 80 kV with a Tecnai G2 electron microscope (FEI, Hillsboro, OR, USA) equipped with a charge-coupled device camera (Advanced Microscopy Techniques, Woburn, MA, USA). Type Ib boutons at M1 were analyzed. For SV counting, T-bars at Ib boutons were identified and Volocity software was used to measure T-bar and PSD length. All data analyses were done blinded.

### MINFLUX Data Acquisition

MINFLUX data were collected on a commercial Abberior Instruments MINFLUX setup (Abberior Instruments GmbH) as previously described^39^. The system was equipped with a 100x/1.4 NA magnification oil immersion lens, a 640 nm continuous-wave laser for exciting the CAGE635-labeled BRP structures in both confocal and MINFLUX mode, a 488 nm pulsed laser for exciting the AF488-co-labeled BRP structures in confocal mode, and a 405 nm continuous-wave laser for photoactivating single CAGE635 molecules. MINFLUX imaging was performed with standard 2D/3D imaging sequences provided by Abberior Instruments and a 640 nm excitation power of 120 μW (2D)/250 μW (3D) in the first iteration, increasing over the subsequent iterations to reach 720 μW (2D)/1.5 mW (3D) in the final one, as measured in front of the microscope body. The emission of individual CAGE635 molecules was detected between 650-720 nm with the pinhole set to 0.83 AU. Before photoactivating CAGE635 during the MINFLUX measurements, confocal images of NMJs were acquired using the AF488 signal of BRP to serve as a guide and identify regions of interest. The sample position was actively stabilized on the back-scattered light of gold beads illuminated with 975 nm in widefield mode. The Abberior Instruments Inspector software with MINFLUX drivers was used to operate the system. MINFLUX data were rendered in a pixel-based fashion.

### Statistical analysis

Statistical analysis and graphing were performed with GraphPad Prism (San Diego, CA, USA). Statistical significance was determined using Student’s t-test for comparisons between two groups, or a one-way ANOVA followed by Tukey’s multiple comparisons test for three or more groups unless noted. In the figures, the center of each distribution is plotted as the mean value and reported in the figure legends as the mean ± SEM with the corresponding number of samples (n). In all cases, n represents the number of individual NMJs analyzed unless noted. The number of larvae (L) used per group in each experiment is indicated in the text. Asterisks in the figures denote p-values of: *, p ≤ 0.05; **, p ≤ 0.01; ***, p ≤ 0.001; and ****, p ≤ 0.0001. The accompanying Table S1 file contains individual spreadsheets labeled with figure number and includes all primary source data and statistical comparisons.

## Data Availability

The authors declare the data supporting the findings of this study are available within the paper, the Source Data file (Table S1) file and the Supplementary Figures. Source data are provided in the paper.

## Acknowledgements

We thank the Bloomington *Drosophila* Stock Center (Indiana University, Bloomington, IN; NIH P40OD018537), the Developmental Studies Hybridoma Bank (University of Iowa, Iowa City, IA), Gerry Rubin, Kate O’Connor-Giles, Ethan Graf, Stephan Sigrist and Dion Dickman for providing *Drosophila* stocks or antisera. We thank Dina Volfson for assistance with strain generation and members of the Littleton lab for helpful discussions. This work was supported by NIH grants MH104536 and NS117588 to J.T.L. and the Freedom Together Foundation.

## Author contributions

Y.A.: Conceptualization, Methodology, Formal Analysis, Investigation, Writing – Original Draft, Visualization; J.M.: Methodology; J.T.L.: Conceptualization, Methodology, Writing – Original Draft, Supervision, Funding Acquisition.

## Supplemental Figures

**Supplemental Fig. 1.**
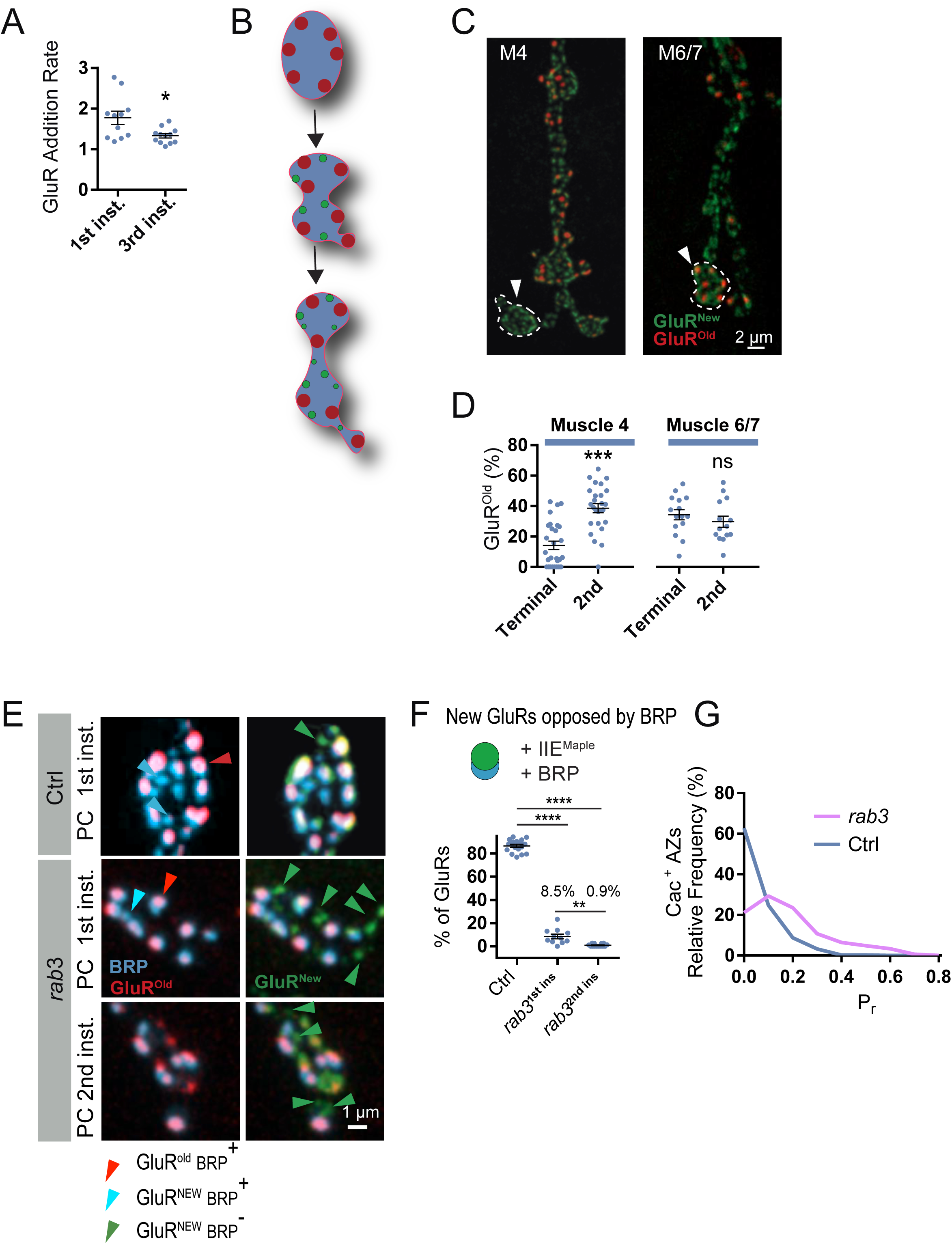
Distinct patterns of synaptic growth at M4 versus M6/7 NMJs and effects of synapse age on AZ maturation in *rab3* mutants. **A.** The ratio of new (green^+^ red^-^ GluR^New^) versus pre-existing (red^+^ GluR^Old^) PSDs was determined 24-48 hours after PC in the 1^st^ instar or 3^rd^ instar stage. The rate of PSD addition during larval NMJ growth is greater during the 1^st^ instar stage compared to 3^rd^ instar (1^st^ instar: 1.78-fold ± 0.16, n=11 NMJs in 4 larvae; 3^rd^ instar: 1.33-fold ± 0.054, n=12 NMJs in 4 larvae, *p* < 0.0133). **B.** Schematic highlighting movement of individual synapses during NMJ growth and new bouton addition. **C.** Representative images of NMJ growth at M4 (left) and M6/7 (right) four days after PC of GluRIIE^Maple^ 1^st^ instar larvae. Note the addition of terminal boutons lacking older red^+^ PSDs (arrow) at M4 versus the internal addition of new boutons lacking red^+^ PSDs at M6 (white outline and arrows). **D.** Quantification of the percent of GluR^Old^ PSDs to all PSDs at the terminal bouton versus the second bouton from the end. Older PSDs are enriched in the terminal bouton at M6/7 NMJs (terminal bouton GluR^Old^ positive PSDs as a percent of all PSDs: M6/7: 34.3% ± 3.4, n=15 NMJs from 5 larvae; M4: 14.2% ± 2.7, n=29 NMJs from 5 larvae; 2^nd^ bouton from end: M6/7: 29.7% ± 3.7, n=14 NMJs from 5 larvae; M4: 38.6% ± 3.0, n=26 NMJs from 5 larvae). **E.** Representative images of 3^rd^ instar M4 boutons immunostained for BRP (cyan) after PC of GluRIIE^Maple^ at the 1^st^ or 2^nd^ instar stage in *rab3* mutants or the 1^st^ instar stage in controls. BRP^+^ AZs were rarely observed at newly formed green^+^/red^-^ GluR^New^ PSDs (green arrowheads) compared to older red^+^ PSDs (red arrowheads) after PC at the 1^st^ instar stage in *rab3* mutants. When PC was performed in the 2^nd^ instar stage in *rab3* mutants, only older red^+^ PSDs were opposed to AZs that were BRP^+^. AZs opposed to newly formed green^+^/red^-^ PSDs were BRP^-^. Controls showed a normal pattern of synapse maturation, with AZs opposed to older red^+^ PSDs containing BRP and 89% of newly formed green^+^/red^-^ PSDs opposed to AZs being BRP^+^. **F.** Quantification of BRP^+^ AZs opposed to newly formed green^+^/red^-^ PSDs after PC in the 1^st^ instar stage in controls and *rab3* mutants (Ctrl: 86.6 ± 1.3%, n=18 NMJs from 3 larvae; *rab3*: 8.5 ± 2.0%, n=11 NMJs from 3 larvae; *p* > 0.001) and after PC in the 2^nd^ instar stage in *rab3* mutants (0.9 ± 0.3%, n=14 NMJs from 3 larvae). **G.** Relative *P_r_* frequency distribution across the AZ population for *rab3* and control M4 NMJs reveals a right-shifted and more homogenous distribution in *rab3* mutants. Statistical significance determined with one-way ANOVA followed by Tukey’s multiple comparisons test. Asterisks denote p-values of: *, p ≤ 0.05; **, p ≤ 0.01; ***, p ≤ 0.001; and ****, p ≤ 0.0001.

**Supplemental Fig. 2.**
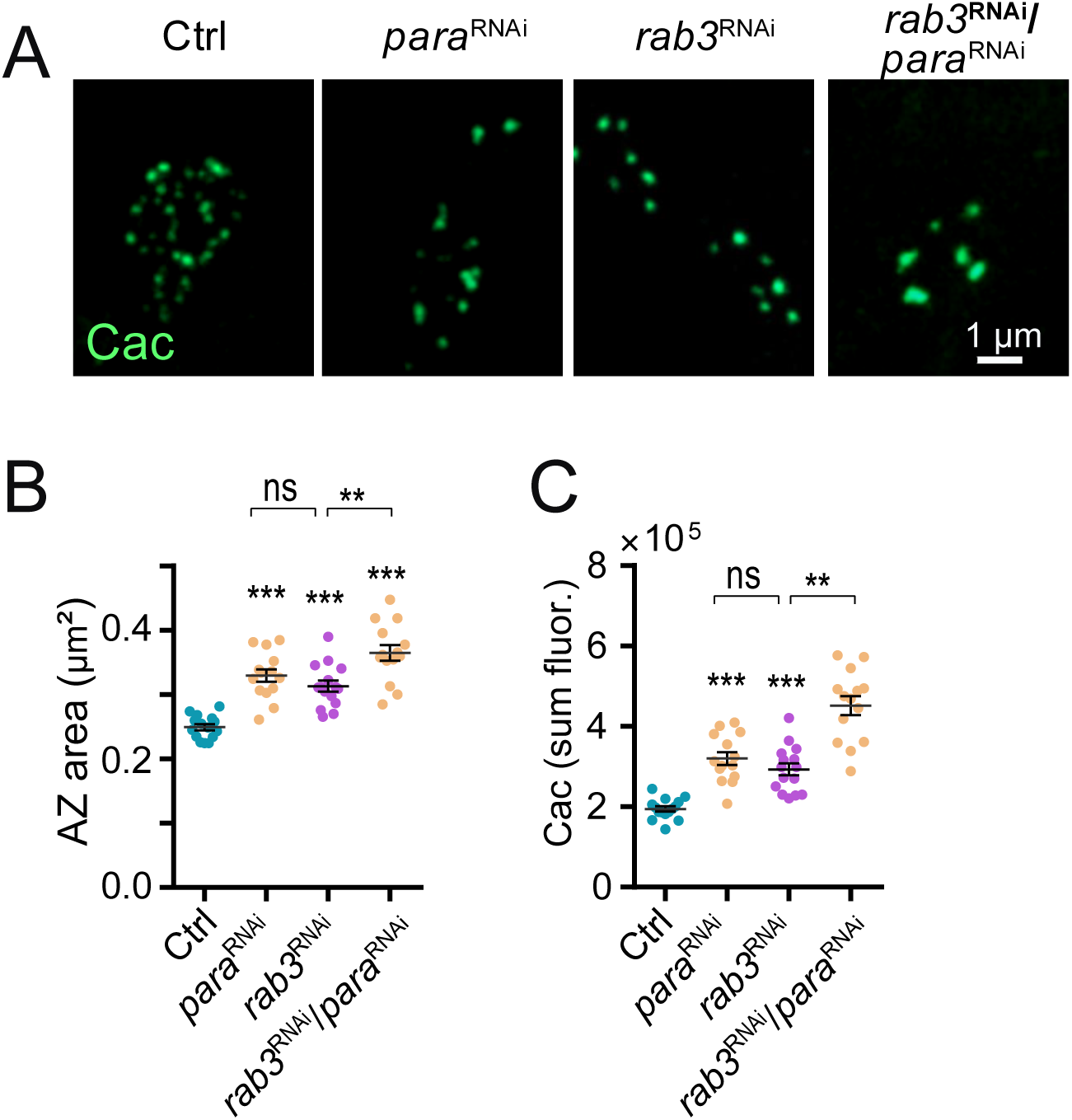
Presynaptic knockdown of the Para sodium channel mimics effects of blocking SV fusion on AZ enlargement. **A.** Representative images of Cac immunolabeling within synaptic boutons of 3^rd^ instar M1 NMJs of controls (left panel) or following expression of *para* RNAi (2^nd^ panel), *rab3* RNAi (3^rd^ panel), or *para* RNAi + *rab3* RNAi (right panel) with the MN1-Ib-Gal4 driver. RNAi-mediated suppression of para also triggers AZ enlargement that is enhanced following Rab3 knockdown. Scale bar, 1 µm. **B.** Quantification of M1 AZ Cac area in the four genotypes (Ctrl: n=15 NMJs from 4 larvae*; para* RNAi: n=14 NMJs from 4 larvae, *p* < 0.0001; *rab3* RNAi: n=15 NMJs from 4 larvae, *p* < 0.0001; *para* RNAi + *rab3* RNAi: n=14 NMJs from 4 larvae, *p* < 0.0001). **C**. Quantification of M1 AZ Cac sum fluorescence in the four genotypes (Ctrl: n=15 NMJs from 4 larvae*; para* RNAi: n=14 NMJs from 4 larvae, *p* < 0.0001; *rab3* RNAi: n=15 NMJs from 4 larvae, *p* = 0.0002; *para* RNAi + *rab3* RNAi: n=14 NMJs from 4 larvae, *p* < 0.0001). Statistical significance determined with Student’s t-test, asterisks denote p-values of: ***, p ≤ 0.001.

